# How does *Mycoplasma pneumoniae* steal lipids from its host membranes?

**DOI:** 10.1101/2023.10.24.563710

**Authors:** Sina Manger, Serena M. Arghittu, Lasse Sprankel, Jakob Meier-Credo, Konstantin Wieland, Daniela Bublak, Julian Langer, Roberto Covino, Achilleas S. Frangakis

**Author notes:** Correspondence: A.S.F. and R.C. These authors contributed equally: Sina Manger, Serena M. Arghittu.

## Abstract

Lipid acquisition and transport are fundamental processes in all organisms. Here, we investigate the lipid uptake and delivery mechanism of the minimal model organism *Mycoplasma pneumoniae*. We show that the essential protein P116 can transport lipids between liposomes independently and without ATP consumption. Our structural data and molecular dynamics simulations reveal the mechanism by which the N-terminal region of P116 perturbs the membrane, the lipid transfer route, and the regulation of membrane binding by the cargo mass within P116’s large hydrophobic cavity. When adequately filled with cargo, P116 undergoes a rapid conformational change that modulates membrane binding. Taken together, our results show that *Mycoplasma* developed one integrated lipid uptake and delivery machinery that simplifies the complex multi-protein pathways used by higher developed organisms.

## Introduction

Biological membranes are essential structures that enclose cells and organize their compartments. Cells carefully regulate their membrane composition (*1–3*). Non-vesicular lipid transport plays a primary role in establishing membrane identity (*4*, *5*).This process relies on diverse protein-based machinery, varying in their components, structure, and mechanisms of action, to transport lipids from one membrane to the other in a controlled way (*6–8*). To effectively understand and control membrane regulation, we must uncover the principles underlying these lipid transport systems.

Lipid transfer proteins (LTPs) range from simple soluble proteins to complex multi-component systems connecting different organelles (*6*).While some LTPs work independently without energy input or additional factors, others require an orchestrated effort from multiple proteins and directly consume energy. Regardless of their complexity, all LTPs must accomplish three fundamental tasks: disrupt thermodynamically stable membranes to extract lipids, transport them across space, and insert them into another stable membrane in a directionally specific manner (*5*, *6*, *9*). Despite their importance, the molecular mechanisms behind these essential steps remain largely unknown (*2*, *6*).

*Mycoplasma pneumoniae*, the primary causative agent of atypical pneumonia in humans, adapts its membrane composition to its host by scavenging host lipids (*10*, *11*). This adaptation serves a dual purpose: It enables opportunistic immune evasion (*12–14*), while fulfilling an essential metabolic requirement, as *M. pneumoniae*’s simplified metabolism cannot synthesise crucial lipids, including phosphatidylcholine, sphingomyelin, and sterols (*10*, *11*).

The membrane-anchored homodimeric protein P116, located on the bacterial surface, plays an essential role in lipid acquisition (*15*). P116 structure comprises two large hydrophobic cavities responsible for lipid binding (*15*). Moreover, as shown by mass spectroscopy analyses, P116 binds a diverse array of lipids, including lipids non-essential for mycoplasma metabolism (*15*). While we have detailed P116’s structure and lipid affinity in a previous study, its transport mechanism and regulation remain unknown.

Here, using an integrated approach combining cryo-electron microscopy, fluorescence-based lipid transfer assays, and molecular dynamics (MD) simulations, we demonstrate that P116 functions as a self-sufficient lipid transfer protein. Our results establish that P116 can extract and transfer lipids independently, without requiring other proteins, cofactors, or direct energy consumption. MD simulations elucidate P116’s membrane binding and lipid transport mechanisms. Our data suggest that the interplay of cargo mass and lipid composition gradient governs lipid transfer, P116’s structural dynamics, and protein-membrane interactions. By revealing how a single protein can autonomously orchestrate complex lipid transport, our findings establish a new paradigm for understanding minimalist biological transport systems and offer opportunities for designing synthetic molecular transporters (*16*, *17*).

## Results

### Comparison with other LTPs suggests that P116’s ectodomain is a functional LTP

Initially, we focused on establishing whether P116 can act as a self-sufficient lipid transfer protein. As established in our previous study, P116 is a homodimer of two 116 kDa monomers, each comprising a core domain, a dimerisation domain, and a N-terminal domain connected to a transmembrane helix via an unstructured linker. The core domain includes four pairs of amphipathic α-helices (the ‘finger helices’), a long α-helix, and a five-strand β-sheet (the ‘palm’), forming a large hydrophobic cavity (Fig. S1a & b).

To create an efficient experimental setup, we simplified our P116 model to include only the essential domains of P116. Therefore, we compared the structure of P116 with other, known self-sufficient transporters. P116’s core domain has a unique fold and contains the lipid pockets. In addition, recent work supports the involvement of its C-terminus in lipid transport (*15*, *18*). In parallel, the N-terminal domain resembles small lipid shuttles like SMP domains, MlaC, and SCP2 (Fig. S2a). Although the amino acid sequences of the P116 N-terminal and E-Syt SMP domains show only 14% sequence identity and 25% similarity with 55% gaps (Fig. S2b), they share a large antiparallel β-sheet surrounding a hydrophobic channel, closed on one side by an α-helix (Fig. S2c). Both SCP2 and SMP domains disrupt membranes, exposing lipids to the aqueous environment, and transfer lipids self-sufficiently (*19*, *20*). This structural similarity of the N-terminal domain of P116 with lipid transfer proteins hints at its involvement in membrane docking. Consequently, we based our further analyses on the ectodomain of P116, which includes the N-terminal domain and the core domain, including its C-terminus, to serve as our model.

### The P116 ectodomain self-sufficiently transfers lipids in vitro

After selecting the model, we established a fluorescence-based assay using liposomes (Fig. 1a) to examine whether P116 can self-sufficiently deposit its cargo into a phospholipid membrane. We depleted the lipid cargo of P116 by detergent treatment and incubated this empty P116 with small (200 nm) fluorescent donor liposomes. We selected lipids for which P116 had shown a higher affinity in our previous study6 and labeled one of the lipid species at the tail and the other at the head with small fluorophores, ensuring that their excitation maxima were sufficiently distinct to enable the use of separate laser lines. The donor liposomes comprised 30% DOPE labeled with a Dansyl-fluorophore at the head group, 30% DOPC labeled with an NBD fluorophore in one acyl chain, and 40% DOPG as a structural scaffold (Fig. 1a & b). We then removed the donor liposomes by two rounds of ultracentrifugation. P116 loaded with fluorescent lipids was subsequently incubated with large acceptor liposomes (> 1 µm) for 2 hours at room temperature. To gain information about the influence on the membrane phase on delivery by P116, we performed the assay using either DOPC (liquid-crystalline phase) or DPPC (gel-phase) acceptor liposomes. In the final step, we removed P116 by four rounds of ultrafiltration at 300,000 MWCO. The remaining acceptor liposomes displayed an intense fluorescence signal (Fig. 1c, Fig. S3). In addition, we included a control accounting for auto-fluorescence of the aceptor liposomes and a control accounting for unspecific transfer, using the non-lipid binding mycoplasma protein P110/P140. The validity of this assay crucially depends on the efficient removal of the fluorescent donor liposomes from P116 and, subsequently, the removal of P116 from the final acceptor liposome sample. Details on the methodology and implemented control experiments can be found in the Material and Methods section.

**Fig. 1.:**
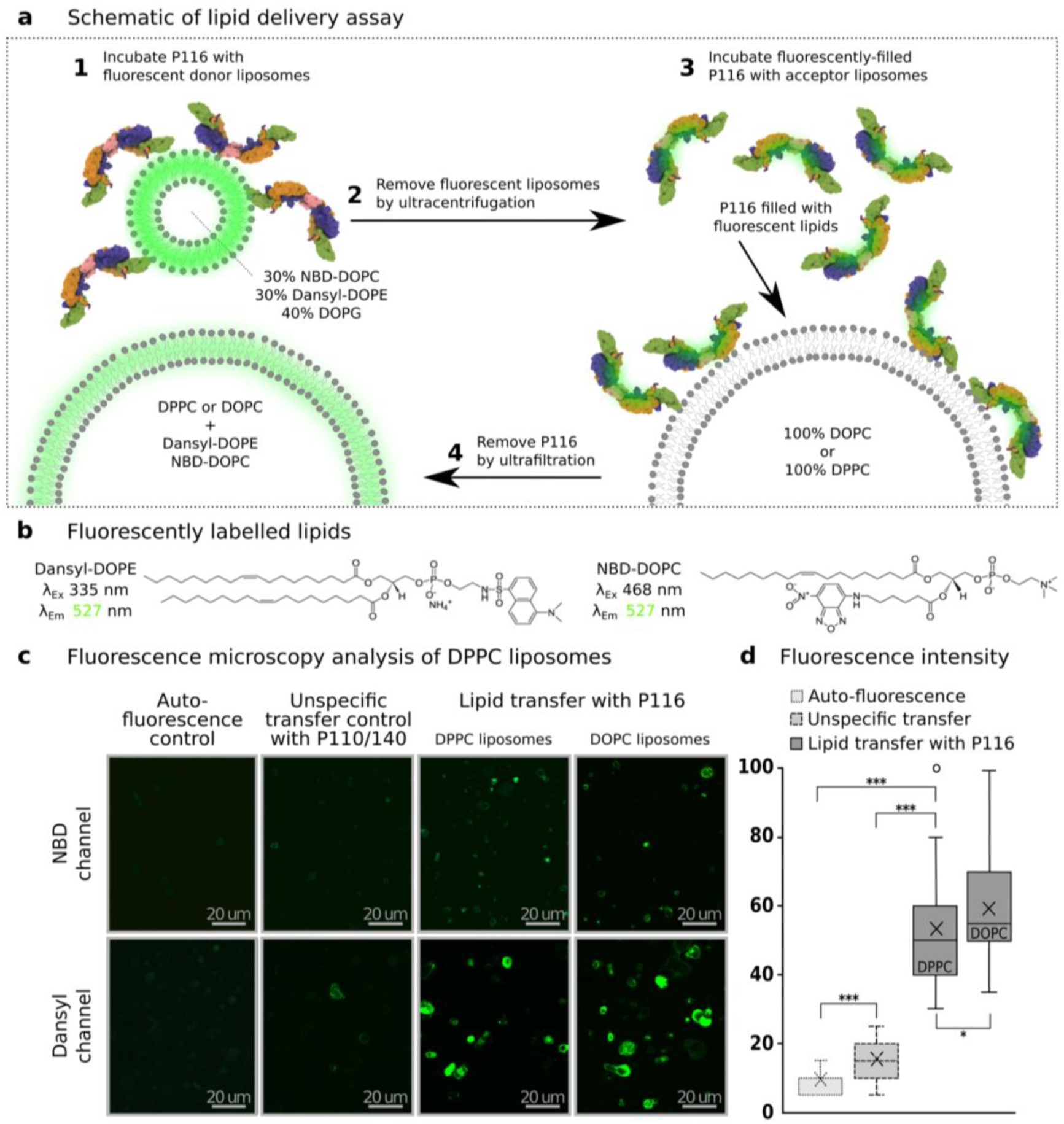
P116 self-sufficiently and single-handedly delivers lipids. (**A**) Schematic of the lipid delivery assay. 1) Incubation of *empty* P116 with fluorescent donor liposomes. 2) Removal of donor liposomes by ultracentrifugation and verification of removal by mass spectrometry. 3) Incubation of P116 filled with fluorescently labelled lipids with non-fluorescent DOPC or DPPC acceptor liposomes. 4) Removal of P116 by ultrafiltration and verification of removal by dot blot, followed by analysis of fluorescent traits of acceptor liposomes by laser-scanning microscopy. (**B**) The chemical structure of the fluorophore labelled lipids (Dansyl-DOPE and NBD-DOPC) used in donor liposomes. Their excitation is at 335 nm and 468 nm, respectively, while their emission is at 527 nm (green spectrum). (**C**) Representative confocal light microscopy images. (Left column) Liposomes (auto-fluorescence control), (middle column) sample from the same workflow with P110/140 instead of P116 (unspecific transfer control), and (right column) sample with P116, according to the workflow detailed in **a** (lipid transfer with P116). The Dansyl and NBD channels are shown separately because of the significant difference in excitation maxima and resulting use of separate laser lines. (**D**) Statistical analysis of fluorescence differences between the sample in the presence of P116 and the controls. All differences are statistically significant. The experiment was carried out in triplicates, and all results could be replicated. The 25–75% data range is contained in the box, the horizontal line represents the median, the vertical line represents the range within 1.5 interquartile range, the X represents the mean, and the O represents outlier. Data can be found in Table S2.

The signal in the auto-fluorescence and unspecific transfer controls was significantly lower than after lipid delivery by P116 (Table S2), indicating successful delivery of NBD-DOPC and Dansyl-DOPE into the acceptor liposomes. In the NBD channel, no signal was detected in either control, while both controls showed some (auto) fluorescence in the Dansyl channel (Fig. 1c). The unspecific transfer control displayed a significantly higher signal compared to the auto-fluorescence control, suggesting either spontaneous transfer of Dansyl-DOPE from the donor liposome or unspecific transfer. Overall, the signal in the Dansyl channel was stronger than in the NBD channel, indicating more efficient transport of Dansyl-DOPE. The delivery into DOPC liposomes was slightly more efficient than into DPPC liposomes, likely due to the increased membrane fluidity of DOPC liposomes at room temperature.

In parallel, we analysed the DPPC acceptor liposomes with mass spectrometry. Our analysis showed relevant amounts of Dansyl-DOPE and NBD-DOPC only in the acceptor liposomes after lipid transfer by P116 and not in the unspecific transfer control (Fig. S4d). This confirms that the ectodomain of P116 can deposit its cargo into a phospholipid membrane without being tethered to the mycoplasma membrane and without requiring other proteins or ATP.

### The conformational dynamics of P116 is enhanced by membrane proximity

Having established that P116’s ectodomain is a self-sufficient LTP, we aimed to clarify the molecular mechanisms behind this process. To achieve this, we examined how P116’s ectodomain interacts with membranes and how these interactions affect its conformations.To access the timescales required by P116 conformational changes, we performed long equilibrium quasi-atomistic molecular dynamics (MD) simulations of the ectodomain (residues 60-868, see Table S1 and Fig. S1a & b) using the MARTINI 3.0 forcefield. MARTINI was developed to reproduce the interactions between lipids and proteins accurately, as documented by numerous validations against experimental data (CIT. # 13 and CIT MARTINI 3 paper). We chose the same membrane composition used in the lipid transfer assay to enable a direct comparison. Our modelling choices and parameters are described in detail in the Material and Methods section.

We simulated *empty* P116 in solution and *full* P116 (9 lipids per cavity) in solution and on the membrane. Our MD simulations showed a broad range of motion involving changes to the *arc diameter* (distance between the monomers), the *wringing angle* (rotation of the monomers with respect to each other), and the *interdomain angle* between the N-terminal domain and the core domain (Fig. S5a).

The MD simulations showed that P116 is highly flexible when close to membranes. To validate the predictions of our MD simulations, we investigated the flexibility of native P116 on the mycoplasma membrane using cryo-ET. Tomography of wild-type mycoplasma cells displays densities at the plasma membrane that resemble P116 (Fig. S6a). To increase the copy number, we generated an *M. pneumoniae* strain that overexpresses *p116* from its native promoter. The tomograms display cells decorated with densities resembling P116 that create a crown-like arrangement on the mycoplasma membrane (Fig. S6a). Due to the visually discernible flexibility of the individual particles (Fig. 2a), we used particle side views and constrained the search range. The resulting density map displayed the core and the N-terminal domain, whose arrangement differed noticeably from the cryo-EM structure while matching a conformation predicted by the simulations (Fig. S1b and S5b). Moreover, classification of the individual particles revealed an extensive range of conformations, many closely resembling those observed in our MD simulations. Cryo-ET confirms that P116 is much more flexible when in the vicinity of membranes and corroborates the accuracy of our P116 molecular model (Fig. 2a, Fig. S6d & c).

**Fig. 2.:**
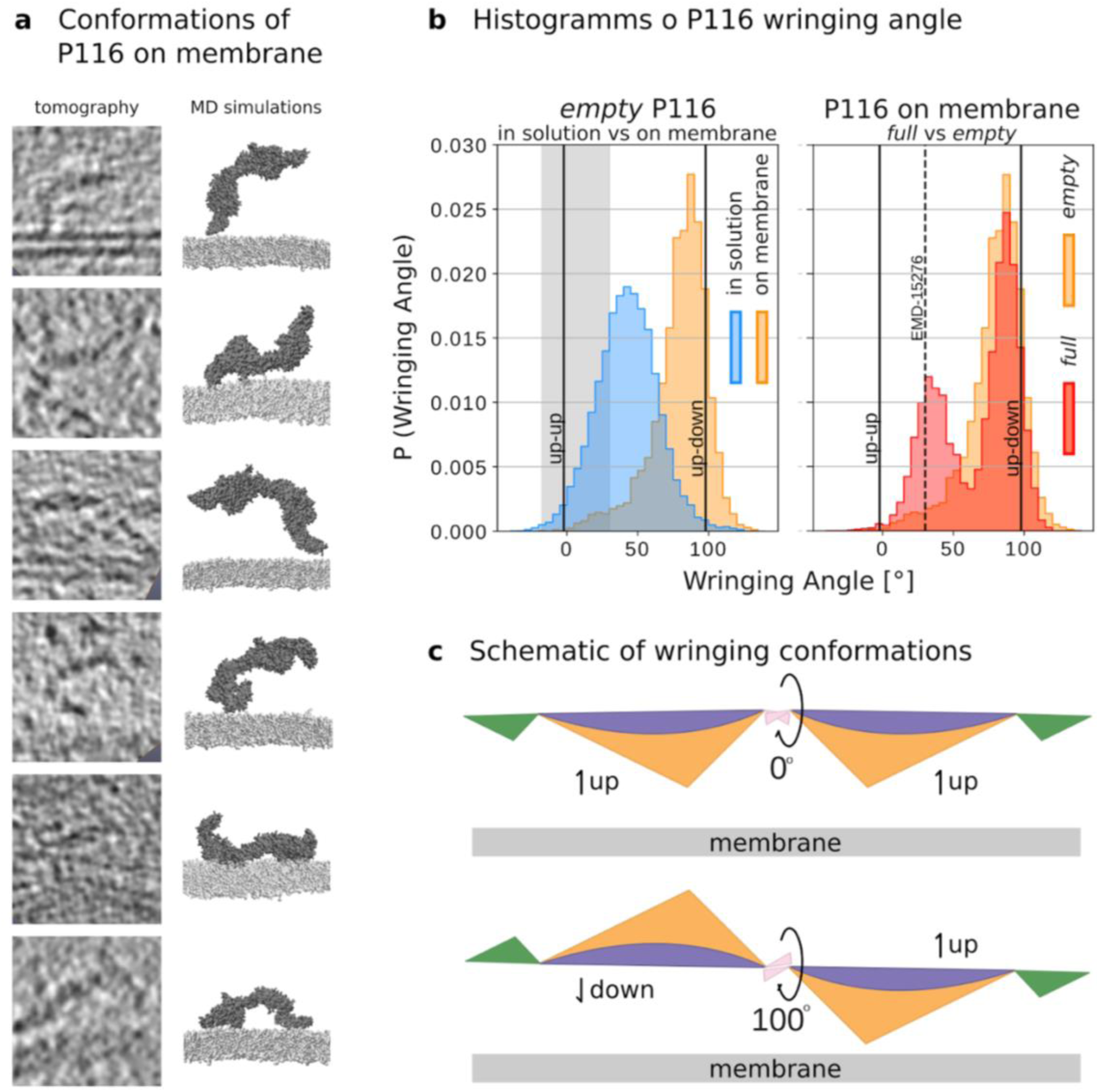
Interaction of P116 with membranes facilitates an increased range of motion, as shown by cryo-ET and MD simulations. (**A**)Gallery of sub-tomographic projection slices of individual particles in various conformations on the membrane is paired with conformations derived from MD simulations. In all simulation renders, P116 (60–868) is represented by a dark gray-filled volume. The membrane is indicated by the phosphate moiety (represented by silver beads). Water and ions are not shown for clarity. (**B**) Histograms of P116 wringing angle from MD simulations. Two graphs are shown: (i) for *empty* P116 in solution and on the membrane and (ii) for *full* vs *empty* P116 on the membrane. (i) Empty P116 adopts a different conformation in solution (blue histogram) compared to when found on the membrane (orange histogram). (ii) The presence of cargo produces two populations of the wringing angle distribution. The cargo-dependent population (shown by the red histogram) at a wringing angle of ∼30° corresponds with the wringing angle observed in the cryo-EM structures of *refilled* P116 (EMD-15276, dashed line). The cargo mass in our *full* P116 model is not as large as in the cryo-EM structure (Fig. S7a). (**C**) Schematic of orientations of the monomers during up-up and up-down wringing conformations. The membrane position is indicated.

### P116’s wringing recapitulates the protein-membrane binding and protein cargo

Among the different motions we observed in the MD trajectories, the wringing angle between the monomers best describes the conformation of P116 (Fig. S5b). In the empty P116 in solution, the wringing angle was 40° 20°, allowing the monomers to face in the same direction (up-up). Conversely, when empty P116 is interacting with the membrane, we observed a wringing angle of 80° 20°, which enables the monomers to face opposite directions (up-down), i.e., one is facing the membrane and the other the medium (Fig. 2b, left & c). The difference in the wringing motion of the protein in solution as compared to the protein in membrane proximity relates to the localization of the membrane binding interfaces of P116 and the protein geometry. We discuss them in the next paragraph. *Full* P116 on a membrane has two wringing angle populations: (i) a wringing angle of 90° 10°, which corresponds to that of empty P116 bound to the membrane and (ii) a second population with a wringing angle of around 30° 10°, which corresponds to the wringing angle observed in the refilled cryo-EM structure (EMD-15276) (Fig. 2b, right). The 30°-population appears directly related to the presence of the cargo.

### MD simulations describe the mechanism of membrane docking

Next, we analyzed how P116’s wringing correlates with membrane binding. During our simulations on a membrane, *empty* P116 recurrently assumed two membrane-bound conformations (Fig. 3a):

i. ‘D-docking’: P116 interacted with the membrane using its up-side, which contains negative charges that interact with lipid head groups (Fig. 3a, *left*, Video S1). During the simulations, D-docking induced membrane curvature (Fig. 3a, *left*).
ii. N-docking’: P116 interacted with the membrane with its down-side and inserted amino acids F860 and F856 into the membrane. Additional insertion of F214 and F227 or F87 further stabilised the docking (Fig. 3a & c). Once stabilised, P116 remained membrane inserted for up to 20 µs and positioned its distal core access (DCA) in close proximity of the membrane (Fig. 3d, Video S1). In all *Mycoplasma* species, residue 214 is a hydrophobic amino acid, and F227 is conserved. In P116, F214 and F227 are part of the N-terminal domain β-sheet, and F87 is part of an N-terminal domain loop. F856 and F860 are situated on the C-terminal phenylalanine-rich loop (the F-loop, 848–868: FAFVDGF) of the core domain, right next to the N-terminal domain (Fig. S1a & b). The C-terminal and N-terminal loops harboring F856, F860, and F87 are part of the hinge region between the N-terminal and core domains.

**Fig. 3.**
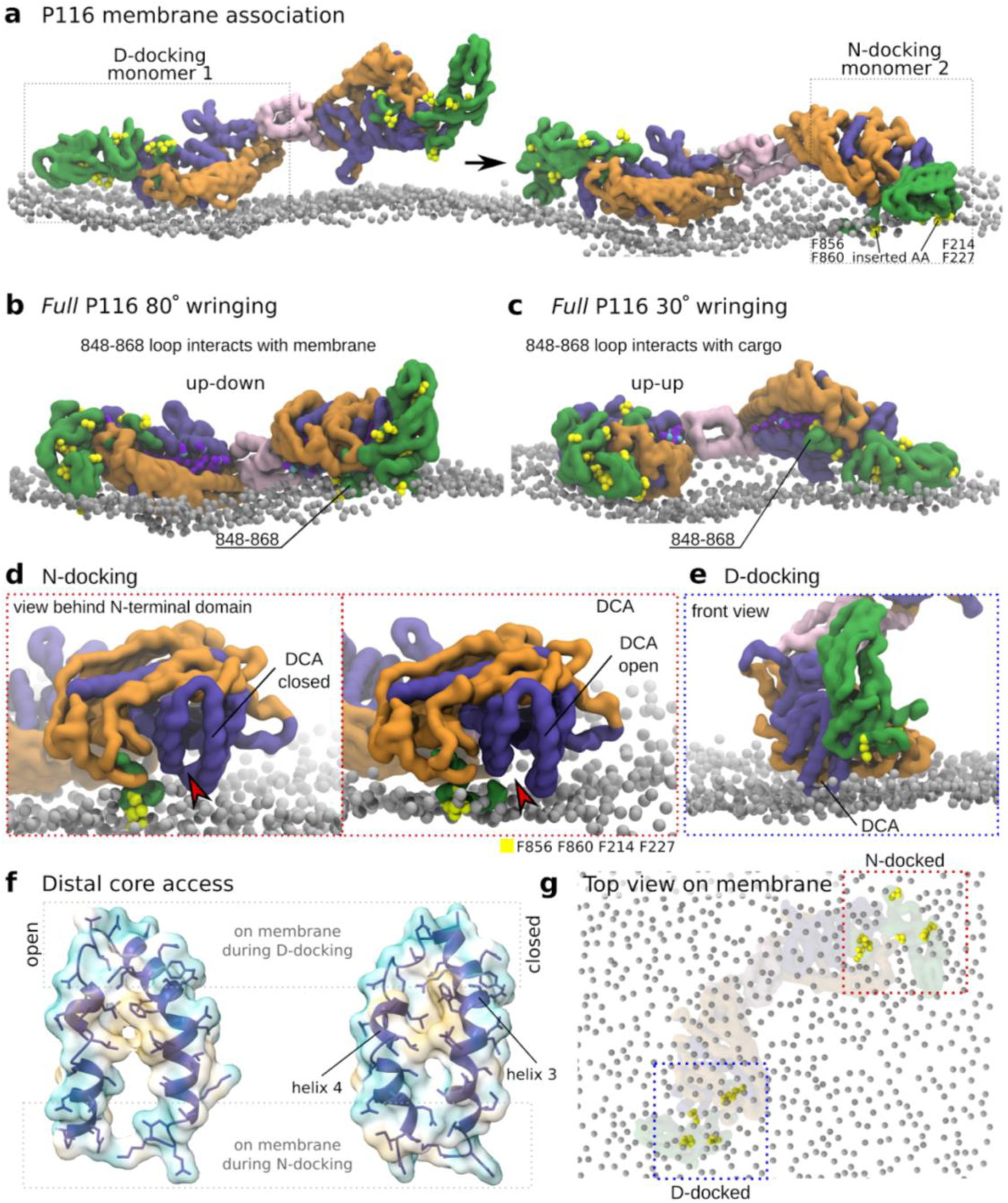
P116 stably docks to membranes in MD simulations. In all simulation renders (**A**-**E**, **G**), P116 (60–868) is represented by a filled volume or cartoon representation colored by domain. When present, the membrane is indicated by the lipid phosphate moieties (represented by silver beads; only one leaflet of the membrane is shown). Water and ions are not shown for clarity. (**A**) P116 renders from MD simulations. The attachment of P116 to a membrane involves phenylalanine residues (yellow beads) in the N-terminal domain. D-docking of monomer 1 precedes N-docking through insertion of F227 and F214 (shown in yellow) of monomer 2. (**B**) MD simulation of *full* P116 at 80° wringing angle. One monomer is N-docked with the F-loop (848-868) and F87 inserted in the membrane, and the other monomer is D-docked. All phenylalanine residues on the N-terminal domain are shown as yellow beads. (**C**) MD simulation of *full* P116 at 30° wringing angle. The F-loop (848-868) interacts with the cargo inside the hydrophobic cavity. No phenylalanine residues are inserted stably into the membrane. All phenylalanine residues on the N-terminal domain are shown as yellow beads. (**D**) N-docking places the DCA near the membrane so that opening (right) and closing (left) happen in direct vicinity to the membrane. Residues F860 and F856, which insert into the membrane, are shown as yellow beads. The N-terminal domain is not shown. (**E**) D-docking places the other side of the DCA near the membrane. Residues F860, F856, and F87 are shown as yellow beads. (**F**) Cryo-EM structures of DCA architecture show opening at one side only. Comparison of the DCA conformation in the *full* and *empty* structures (PDB: 8A9A and 8A9B, respectively). The two helices comprising the DCA are shown in blue with the individual amino acid sidechains as stick models. The surface representations are colored by the hydrophobicity (yellow is hydrophobic, and blue is hydrophilic). The DCA opens and closes at the side that is facing the membrane during N-docking, while the other side remains closed. (**G**) N-docking facilitates the spread of the membrane beneath the DCA. Top view of the membrane during docking of P116. The D-docking area is labelled with a blue box and the N-docking area with a red box. Residues F214, F227, F856 and F860 appear as yellow beads. The membrane structure is disrupted only during N-docking, which facilitates the spreading of the membrane beneath the DCA.

In the simulations, we observed a consistent sequence of events: initially, one monomer engaged with the membrane in a D-docking conformation, followed by the N-docking of the second monomer (see Fig. 3a), which stabilised the binding. This sequence of events suggests that D-docking might be involved in membrane recognition. In parallel, when P116 is N-docked to the membrane, the interdomain angle range of the two monomers differs (Fig. S5c). While the monomer attached to the membrane has a fixed configuration of the N-terminal domain, the N-terminal domain of the D-docked monomer can move freely. The confinement of the N-terminal domain during docking, along with its structural similarity to SMP domains, supports its crucial role in stabilising membrane docking in conjunction with the F-loop insertion under the lipid headgroups.

### The cargo mass modulates membrane binding

Following the observation that the wringing angle is affected by the protein’s cargo, we further quantified their correlation. The wringing angle population of 80° for P116 on the membrane (Fig. 2b & 3b) corresponds to simultaneous D- and N-docking. In contrast, the 30° wringing angle population represents a conformation where N-docking does not occur. In this conformation, the phenylalanines on the terminal F-loop interact with the cargo inside the cavity instead of inserting into the membrane, thereby preventing N-docking (Fig. 2b & 3c). These results suggest that the cargo mass controls the propensity of P116 detaching from the membrane.

To probe this, we simulated four replicas of an intermediate model between the *full* P116 and empty P116 models, having cargo mass reduced by 33% (from 9 to 6 lipids per cavity). We then compared the ratio of the 30° wringing angle population at the different cargo masses. We used a 30° wringing angle configuration from the *full* P116 model as the initial conformation, allowing us to assess the stability of membrane binding when N-docking is hindered. In all four simulations, P116 completely detached from the membrane, indicating unstable binding. Moreover, the partially filled P116 exhibited an intermediate 30° wringing angle population between the *empty* and *full* P116 models (Fig. S7c). These findings suggest that the cargo mass directly modulates membrane binding.

### DCA opening occurs in N-docking but not in D-docking

Although both N-docking and D-docking position the DCA near the membrane (Fig. 3d & 3e), only N-docking allows the DCA to open and close directly adjacent to the membrane. This behavior is due to distance changes between helices 3 and 4, similar to those observed in the cryo-EM structures (Fig. 3d & 3f). Additionally, D-docking induces membrane curvature (Fig. 3a), causing lipid head groups beneath the N-docked monomer to move apart (Fig. 3g), which may facilitate lipid transfer by exposing the membrane core.

### The N-terminal domain and the F-loop are necessary for lipid uptake from intact membranes

Our MD simulation highlighted that both the N-terminal domain and the C-terminal F-loop are crucial for lipid uptake from membranes. To test this hypothesis, we decided to construct a ΔN-P116 variant (residues 246–818; see Table S1 and Fig. S1a & b) with truncated N-terminal domain and F-loop, following the finding that point mutations in the F-loop residues F854, F856, and F860 compromised P116’s structural integrity (Fig. S8).

We determined the structure of ΔN-P116 by single-particle cryo-electron microscopy (cryo-EM) at 4.3 Å (EMD-50314, PDB: 9FCH; Fig. S9a & S10a, Table S3b & S3c). The structure resembled the full-length ectodomain of P116 (residues 30–957; EMD-15274, PDB: 8A9A) without the N-terminal domain. The core domain was filled with lipids within its hydrophobic cavity, as P116 is exposed to various lipids during the expression and purification procedure. The preservation of the protein fold and lipid binding by the ΔN-P116 construct made it a reliable system for further analysis.

Comparing the lipid extraction ability of the full-length ectodomain of P116 and ΔN-P116 poses a technical challenge. P116 contains multiple lipid binding sites, including the central cavity, the channel in the N-terminal domain, and nonspecific binding sites for micelles at the hinge region. Additionally, the central cavities can accommodate up to 20 lipid molecules, which are subject to constant reorientation within the cavity (*15*). Interpreting signals from these interactions using traditional techniques like ITC, MST, or SPR can be misleading (*21*). Therefore, we utilised mass spectrometry to identify the types of lipids present in the sample, along with cryo-EM, to ascertain the localisation of these lipids within the P116 structure.

We first reduced the lipid cargo of P116 by detergent treatment and verified the absence of DPPC via mass spectrometry (Fig. 4a). We additionally confirmed that P116 had assumed the *empty* conformation by determining its structure with cryo-EM at 4.3 Å (Fig. 1b & c, EMD-51408; Fig. S11a, Table S3a). This is possible because the volume of the hydrophobic cavity in the *empty* protein is reduced as helix pair 2 moves inward in the absence of lipid cargo (*15*). Thus, the position of helix pair 2 distinguishes whether the cavity is full or empty (Fig. S12, Table S4). In addition, the cavity in the full state typically contains densities from lipid molecules (Fig. 4e & f). Thereafter, we incubated the *empty* P116 with DPPC liposomes, which we then removed by two rounds of ultracentrifugation. Analysis of the cargo of P116 by mass spectrometry revealed the presence of DPPC, which was previously absent in the *empty* P116 (Fig. 4d). We confirmed the localisation of lipids within the cavity of P116 by determining its structure with cryo-EM at 3.6 Å (Fig. 4e & f, EMD-51409; Fig. S11b, Table S3a). As expected, the soluble ectodomain of P116 can disrupt the membrane and extract lipids on its own.

**Fig. 4.**
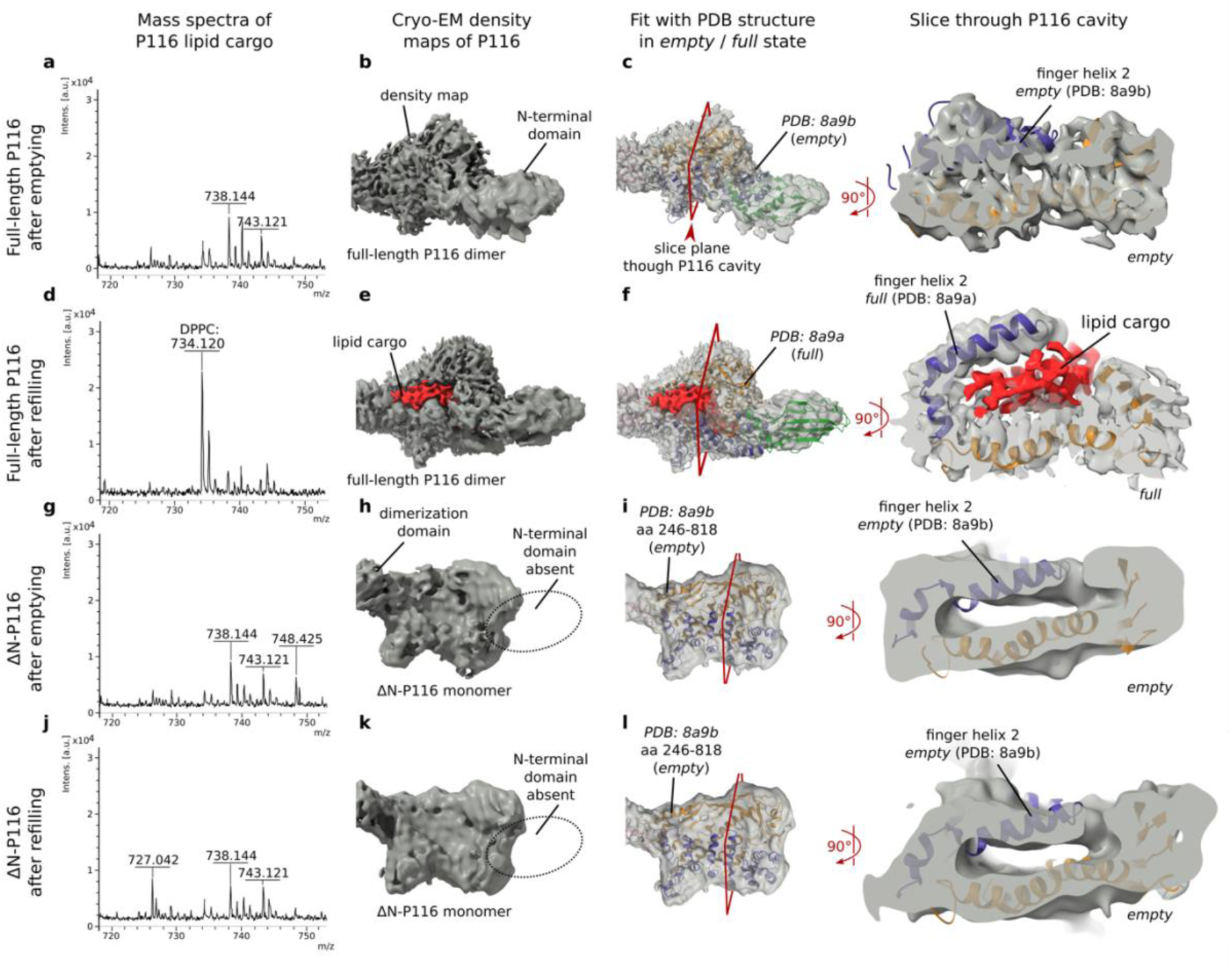
The N-terminal domain of P116 is required to extract lipids from membranes. The 1st column reports mass spectrometry analysis of P116 lipid cargo. The 2nd column reports the isosurface representation of cryo-EM density maps of P116 in dark grey. The 3rd and 4th column report the cryo-EM density maps of P116 superimposed with ribbon models of the *full* (PDB: 8A9A) and *empty* (PDB: 8A9B) conformation of P116, respectively. Correlation values for the respective fits can be found in Table S4. (left) Top view on one monomer. (Right) Computational slice through the cavity of P116. The position of the computational slice with regard to the density map is indicated with red frames. The volume of the cavity determines the conformation, the position of helix pair 2, and the presence and absence of lipids (Fig. S12a & b). (**A**) *Empty* full-length P116 (traces of lipids at the masses 738 and 743 Da are visible). (**B**) *Empty* full-length P116 at 3.6 Å resolution (EMD-51408, Fig. S11b). (**C**) *Empty* full-length P116 superimposed with the ribbon model of the *empty* (PDB: 8A9B) conformation. (**D**) Full-length P116 after incubation with and removal of DPPC liposomes (a clear peak at 734 Da, the mass of DPPC, is visible). (**E**) Full-length P116 after incubation with and removal of DPPC liposomes at 4.36 Å resolution (EMD-51409, Fig. S11a). Lipid densities inside the cavity are shown in red. (**F**) Full-length P116 after incubation with and removal of DPPC liposomes superimposed with the ribbon model of the *full* (PDB: 8A9A) conformation. Lipid densities inside the cavity are shown in red. (**G**) *Empty* ΔN-P116 (traces of lipids at the masses 738, 743, and 748 Da are visible). (**H**) *Empty* ΔN-P116 at 5.3 Å resolution (EMD-51412, Fig. S10b). (**I**) *Empty* ΔN-P116 superimposed with the ribbon model of the *empty* (PDB: 8A9B, truncated to aa 246-818) conformation. (**J**) ΔN-P116 after incubation with and removal of DPPC liposomes (traces of lipids at the masses 727, 738, and 743 Da are visible). (**K**) ΔN-P116 after incubation with and removal of DPPC liposomes at 5.43 Å resolution (EMD-51411, Fig. S10c). (**L**) ΔN-P116 after incubation with and removal of DPPC liposomes superimposed with the ribbon model of the *empty* (PDB: 8A9B, truncated to aa 246-818) conformation.

Next, we emptied the ΔN-P116 and incubated it with DPPC liposomes, following the identical procedure as with the full-length ectodomain. We again determined the presence of lipid cargo after emptying and after attempted refilling by both mass spectrometry and single-particle cryo-EM (Fig. 4g - l, EMD-51412, EMD-51411, Fig. S10b & c, Table S3b & S3c). Unlike the full-length ectodomain, the ΔN-ectodomain did not refill during incubation with DPPC liposomes and instead remained empty (Fig. S12 and Table S4).

Thus, the absence of the N-terminal domain and F-loop dramatically reduces, or completely disrupts, the protein’s ability to extract lipids from intact membranes. Notably, these results agree with preliminary results, where a monoclonal antibody against the C-terminal loop reduced the uptake-ability of P116 and inhibited *Mycoplasma* growth (*18*).

### P116 takes up a lipid molecule via the DCA

Next, we focused on detailing the mechanism through which lipids enter P116. This entails identifying the lipid transfer route and potential intermediate states in the process. We first concentrated on the transfer route. In *empty* P116, the cavity collapses to a fraction of its volume and is only accessible via the DCA or the finger helices (Fig. 5a). To determine which of these two possible access points mediates lipid entry, we performed MD simulations of *empty* P116 with free DOPE and DOPG lipids in solution (Fig. 5b, *left*). This far-from-equilibrium initial condition enabled us to sample the interactions between P116 and the lipids efficiently. The simulations revealed four events: (i) the formation of micelle-like lipid aggregates; (ii) the interaction of the micelle with P116 at N-terminal domain tip and hinge region (854–860); (iii) the insertion of a lipid molecule in the channel within the N-terminal domain; (iv) and the interaction of a lipid molecule at the hinge region and subsequent lipid uptake through the DCA of one of the monomers. Of note, the DCAs were closed at the beginning of the simulation.

**Fig. 5.**
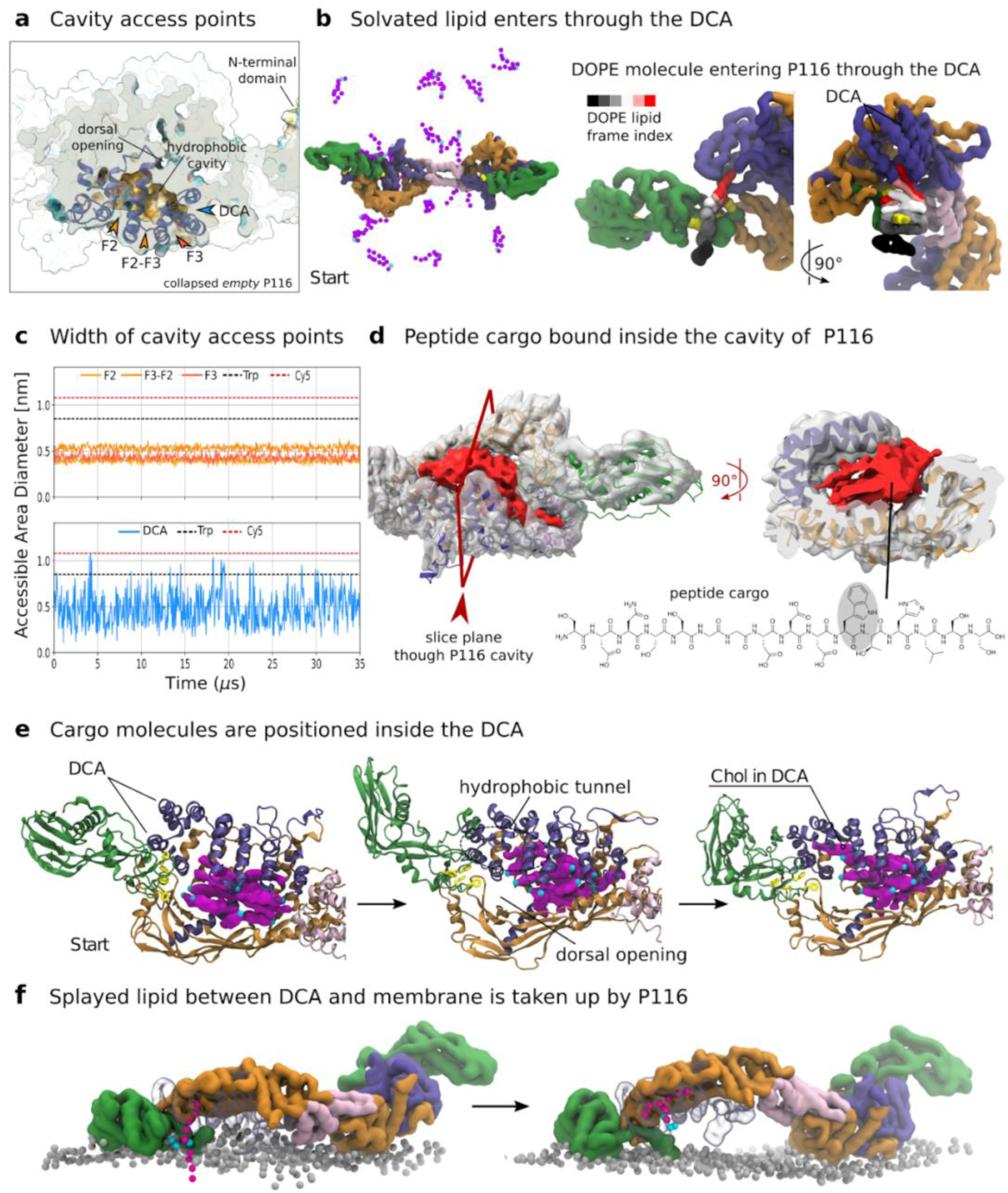
Lipid uptake and delivery occur through the distal core access. In all simulation renders, P116 is represented by a filled volume or cartoon representation colored by domain. When present, the membrane is indicated by the phosphate moiety (silver beads), and solvated lipids are represented by phosphate moieties (blue beads) and tails (purple beads). Water and ions are not shown for clarity. (**A**) Computational slice through the central cavity of *empty* and collapsed P116. The positions of the dorsal opening, the DCA, and the finger helices (F2 and F3) are indicated. The section surface is shown in gray. (**B**) Solvated lipids enter P116 through the DCA. Coarse-grained MD simulation of P116 with free-floating DOPE, DOPC, and DOPG lipids. F860, F856, F227, and F214 are shown as yellow beads. (Left) Start of simulation with evenly distributed lipids. (Right) DOPE lipid entering the cavity through the DCA. The lipid is shown as a filled volume colored according to the trajectory frame. (**C**) Opening width of finger helices and DCA of *empty* P116 measured as accessible surface area throughout 35 µs. (**D**) Cryo-EM density map of P116 superimposed with a ribbon model of the *full* structure (PDB: 8A9A) of P116. *Right,* top view on one monomer. *Left,* computational slice through the cavity of P116. The position of the computational slice with regard to the density map is indicated with red frames. The hydrophobic cavity contains the peptide shown below. Corresponding mass spectra and details on the cryo-EM processing can be found in Fig. S14. (**E**) Cargo inside the cavity is positioned inside the DCA. Atomistic MD simulation of P116’s cavity filled with 12 cholesterol molecules (shown as purple spheres). From left to right: start of the simulation with cholesterol molecules evenly distributed inside the cavity; cholesterol molecules rearrange and are squeezed into the cavity towards the dimerisation domain; and an individual cholesterol molecule inserts into the hydrophobic channel formed by the DCA. (**F**) Lipids are taken up by P116 when placed in a splayed configuration between the membrane and DCA. (Left) N-docked P116 with a splayed DOPE lipid positioned below the DCA. (Right) DOPE lipid entering the cavity through the DCA. The DCA and amphipathic helices of the N-docked monomer are rendered as transparent shapes for clarity.

Prior to uptake, the lipid was first caged at the hinge region outside of the DCA by F854, F856, F860, and F87 (Fig. 5b, *middle*; Video S2). Notably, residues F860 and F856 were inserted into the membrane during N-docking (Fig. 3a). The lipid uptake was then initiated by the splaying of one lipid tail into the DCA, followed by the other. The hydrophilic head group remained on the outside of the cavity during the whole process (Fig. 5b, *right*, Video S2). Once the lipid entered the core cavity, it positioned its head group alternatively towards the solvent-accessible areas at the dorsal opening or below the finger helices. In parallel, the interaction of lipid micelles with the N-terminal domains and the hinge region further corroborates their involvement in membrane association and lipid binding (Video S2). These interactions involved the same phenylalanine residues that were inserted during N-docking on the membrane (F227 and F214, Fig. 4a). Crucially, neither the individual free lipids nor the later-formed micelle interacted with the rest of the protein structure. This behaviour suggests that lipids likely enter P116 through the distal core access. To test the model’s validity, we repeated the same simulation but restrained the DCA to keep it sealed. As anticipated, during this simulation, no lipids entered through the DCA in the cavity, and we could recapitulate all the other interactions.

### The flexibility at the hinge region is necessary for lipid uptake

To investigate the role of the hinge region in lipid uptake, we repeated the simulation of P116 and free-lipids in solution with the protein’s flexibility restricted to the range measured in cryo-EM structures. Despite the spontaneous localisation of a few lipid molecules at the hinge region (Fig. S13b & c), no uptake occurred during the simulation. The misarrangement of the F-loop and the N-terminal domain at the hinge region is responsible for the abolition of lipid uptake, as it hinders the lipid caging prior to lipid splaying and uptake. This implies that lipid caging and membrane insertion required a high flexibility at the hinge region and, consequently, of the F-loop and N-terminal domain, as observed in the cryo-ET dataset (Fig. S6b & c). Crucially, this finding explains our assay showing that both the N-terminal domain and the F-loop are essential for lipid extraction. Additionally, it suggests that the lipid caging at the hinge region is a crucial step of the uptake mechanism prior to lipid splaying.

### The DCA is the only entry point of P116

To verify that cargo uptake happens through the DCA, we conducted binding assays with hydrophobic peptides. Following the measurement of the fluctuating opening diameter of both the finger helices and the DCA, we predicted that P116 should be able to uptake a hydrophobic compound with a maximum diameter below 0.9 nm (∼tryptophan residue), which would fit the DCA but not the finger helices. The same compound labelled with a Cy5-fluorophore (1.1 nm) should exceed the opening diameter of the DCA and thus not be taken up by P116. We incubated P116 with both compounds individually and, after thorough washing, measured lipid binding using mass spectrometry (Fig. S14a). As predicted, P116 took up only the plain peptide. To confirm the peptide was bound within the hydrophobic cavity, we determined the structure of the peptide-filled P116 using cryoEM at 3.3 Å (Fig. 5e, EMD-18476; Fig. S14b & c, Table S5).

### The atomistic simulation of P116 shows the insertion of cholesterol cargo in the DCA

To further study the behavior of the cargo inside the hydrophobic cavity of P116, we performed an atomistic MD simulation of P116 filled with 12 cholesterol molecules per monomer. We used atomistic resolution to optimise for the accuracy of the chemical interactions between cholesterol and the P116 cavity. The amphipathic helices (Helices 2 & 3, Fig. 3f) squeezed the cargo into the cavity towards the center of the dimer. The hydrophobic cavity core remained connected to the DCA through a short narrow tunnel passing over the dorsal opening (Fig. 5e).

During the simulation, cholesterol molecules moved from inside the cavity to the DCA via the connecting tunnel, orienting their hydroxyl group toward the solvent. As the simulation progressed, these cholesterol molecules returned to the cavity and were replaced by other cholesterol molecules from the bulk (Fig. 5e). This behavior of the cholesterol cargo completes our description of the P116’s cavity-lipid interactions and demonstrates that both the delivery and uptake routes pass through the DCA.

### Lipid splaying is the rate-limiting step in P116 lipid extraction from membranes

Having identified the lipid transfer route, we focused on detailing the extraction mechanism on membranes. According to our simulations with lipids in solution, lipid transfer would involve a lipid splaying event. Lipid splaying consists of the spreading of the lipid hydrophobic tails and is a known intermediate state in membrane fusion (*22*, *23*) and lipid acquisition by LTPs (*24*). At the typical timescales of coarse-grained simulations, lipid splaying from a membrane is an extremely rare event and we never observed it. Thus, to test whether lipid uptake is consequential to lipid splaying, we began our simulations with a splayed lipid already positioned between the membrane and the open DCA of an N-docked P116 (Fig. 5f). This setup aimed to reduce the simulation time needed to observe lipid transfer by bypassing the rate-limiting step of lipid splaying (*25*). We ran 48 unbiased replicas of the system for about 14 µs each and observed spontaneous lipid uptake in 40 out of 48 replicas (Fig. 5f, Fig. S13a). We did not apply force to extract the lipid from the membrane; instead, after preparing it in a splayed configuration, uptake happened spontaneously over approximately 10 µs. This extended uptake and time indicates that the lipid was stably splayed between the membrane’s hydrophobic core and the P116 cavity. Together with the collected statistics, this implies that lipid uptake is highly favoured once splaying has occurred.

## Discussion

*Mycoplasma pneumoniae* employs the membrane-anchored protein P116 to adjust its membrane composition to the host by scavenging essential lipids (*15*). Through a fluorescence-based assay, we showed that the isolated ectodomain of P116 acts as a self-sufficient LTP. This finding is particularly significant for synthetic biology, where minimalistic, adaptive lipid transport systems are essential for engineering lipid metabolism in synthetic organisms (*17*).

Integrating computation and experiments, we characterised the transport mechanism of P116. Membrane docking occurs through two coordinated mechanisms: D-docking initiates membrane interaction, and subsequent N-docking stabilises binding by inserting phenylalanine residues under the membrane surface. Biochemical assays and cryo-EM analyses confirmed the critical role of the hinge region in these processes, as predicted by our molecular model. Moreover, molecular dynamics simulations highlighted lipid splaying as an essential precursor to lipid entry and indicated the DCA as the lipid transfer route, a mechanism experimentally corroborated through targeted binding assays using hydrophobic peptide probes for entry size.

Additionally, our simulations revealed a unique cargo-mass-dependent mechanism in which the mass of the lipid cargo modulates P116’s binding stability to the membrane. As lipid cargo accumulates, the interaction between cargo and the F-loop inhibits stable docking, leading to detachment from the membrane upon saturation (Fig. 6). Combined with its large hydrophobic cavities and dimeric architecture, this distinctive mass-based regulation sets P116 apart from other known LTPs, which are typically specialised for specific lipid classes and rely on smaller lipid binding domains (*6*). In contrast, P116 demonstrates remarkable functional versatility, binding a wide range of lipid species (*15*), which is crucial for the adaptability of *Mycoplasma* to diverse host environments. The lipid transport directionality is ultimately ensured by the host, which consumes energy to maintain its outer membrane composition (*2*, *4*, *5*).

**Fig. 6.**
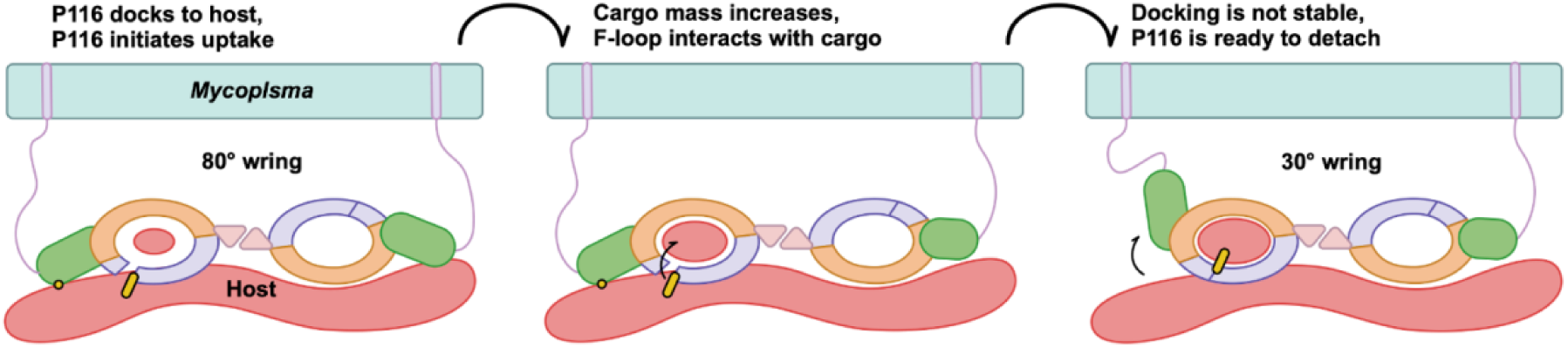
Schematic representation of the self-regulation of P116 via cargo mass. P116’s domains are colored as usual, the F-loop is represented in yellow with black contours, the mycoplasma linker is represented in lilac, the host membrane/lipids is red, and the mycoplasma membrane is oil green. At the beginning of the uptake, the F-loop is anchored to the membrane. When the cargo is sufficiently large, the F-loop likely interacts with it. Without the anchoring of the F-loop, N-docking is not possible, and the protein will eventually detach from the membrane.

While our study emphasises the ectodomain’s autonomous lipid transport mechanism, future research should explore the potential impact of P116’s upper charged interface on its membrane affinity. Different membrane compositions may influence the transport by altering binding affinity and membrane tension. In our simulations, we found that D-docking initiates binding and induces membrane curvature, exposing the hydrophobic core under the N-docked monomer (Fig. 3g). We speculate that reduced host membrane tension may enhance the induced curvature, facilitating lipid exchange. At the same time, although affinity for specific membrane compositions is not essential for P116 transport, it may enhance efficiency by favouring D-docking.

Moreover, further investigation may focus on the transfer of cholesterol between membranes. The proposed lipid transfer mechanism involves lipid splaying, which is not feasible for cholesterol due to its unique geometry. Instead, we speculate that cholesterol uptake may be facilitated by its non-polarity (*26*, *27*). This could lead to cholesterol accumulation beneath the DCA, where the lipid headgroups spread apart due to the hydrophobic effect.

In summary, our findings establish P116 as a minimalist model of lipid transport systems. Elucidating its unique self-sufficient transport mechanism deepens our understanding of lipid homeostasis in minimal organisms and holds substantial promise for the rational design of synthetic lipid transporters with wide-ranging biotechnological applications.

## Materials and Methods

### Expression and purification of P116

The extracellular domain of P116, C-terminally shortened and HIS-tagged (30–957), was expressed via the vector pOPINE_P116 (backbone #26043; Addgene, Watertown, USA). The vector is available upon request. *E. coli* BL21 (DE3) cells harboring the vector were grown at 37°C to an OD_600_ of 0.6, induced with 0.6 mM IPTG, and further grown overnight at 20°C with mild shaking. Cells were harvested, lysed by sonication in lysis buffer (50 mM Tris-HCl, pH 7.4, 20 mM imidazole, 1 mM PMSF) and centrifuged at 20,000 x g in a tabletop centrifuge for 45 min at 4°C. The supernatant was loaded onto a HisTrap 5 mL column (GE Healthcare, Chicago, USA) that was pre-equilibrated in binding buffer (20 mM Tris-HCl, pH 7.4, 20 mM imidazole), thoroughly washed in binding buffer and eluted with elution buffer (20 mM Tris-HCl pH 7.4, 400 mM imidazole, 150 mM NaCl). The eluate was loaded onto a Superose 6 column (GE Healthcare, Chicago, USA) in protein buffer (20 mM TRIS-HCl, pH 7.4, 150 mM NaCl). Fractions containing non-aggregated P116 were pooled and stored at –80 °C.

### Emptying of P116

To obtain *empty* P116, we added 2.6% Triton X-100 to the protein sample and incubated it for either 1.5 h at room temperature (full-length P116) or 3 h at 37°C (ΔN-P116) with shaking. The Triton X-100 was then removed by either (i) binding to a HisTrap 1 mL column (GE Healthcare, Chicago, USA) or HIS-Select Nickel Affinity Gel in a gravity-flow column (Merck, Darmstadt, Germany), followed by washing with binding buffer with and without 1.3% Triton X-100, before eluting the samples from the column; or (ii) diluting twice to 15 mL and then to 95% concentration in a centrifugal concentrator (Pierce, 50,000 MWCO PES; Thermo Fischer Scientific, Waltham, USA) and subsequently running through a detergent removal spin column (Pierce; Thermo Fischer Scientific, Waltham, USA). P116 was concentrated using Vivaspin 500 centrifugal concentrators (10,000 MWCO PES; Sartorius, Göttingen, Germany) to a final concentration of >0.5 mg/mL.

### Preparation of liposomes

Generally, lipids were dissolved in chloroform, then mixed and dried under nitrogen followed by high vacuum overnight. The dried lipid mixtures were then slowly dispersed in liposome buffer (20 mM Tris-HCl, pH 7.4) and left to hydrate for 30 min with gentle agitation. The liposomes were extruded with an Avanti Mini-Extruder (Avanti Polar Lipids, Alabaster, USA) and stored at 4°C in the dark.

Fluorescent liposomes (30% Dansyl-DOPE, 30% NBD-DOPC and 40% DOPG) were prepared from 1 mg of Dansyl-DOPE, 0.75 mg NBD-DOPC and 1 mg DOPG, respectively (Avanti Polar Lipids #810330c, #810132c and #840475c, respectively). Hydration and extrusion were performed at room temperature to a final concentration of 2.75 mg/mL and a diameter of 200 nm. DPPC and DOPC liposomes were prepared with the pure lipids (Avanti Polar Lipids DPPC #850355c, DOPC # 850375c, respectively). Hydration was carried out at 50°C and 25°C, respectively, to a final concentration of 3 mg/mL. The liposomes were not extruded.

### Lipid transfer assay

To fill P116 (30 – 975) with fluorescent lipids, 0.5 mg of emptied P116 was incubated with 50 µL of fluorescent donor liposomes (30% Dansyl-DOPE, 30% NBD-DOPC, 40% DOPG) for 2 h at 37°C with gentle shaking. Before the experiment, the donor liposomes were freshly extruded through a 200 nm filter, ensuring that their size was below the resolution limit of a laser scanning microscope with a 63x/1.4 oil objective.

The fluorescent liposomes were then removed by ultracentrifugation for 30 min at 150,000 x g in a Ti70 rotor with 1.5 mL Eppendorf tube adapters and re-ultracentrifugation of the upper 80% of supernatant before collecting the upper 90% of supernatant. Efficient removal was probed by MALDI-TOF mass spectrometry of a replicate experiment, in which the ectodomain of the non-lipid-binding mycoplasma membrane protein P110/P140 was used instead of P116 (Fig. S4a), yielding no detectable DOPE, DOPC or DOPG in the final sample. The supernatant was concentrated using 10,000 MWCO centrifugal concentrators (Vivaspin 500, PES; Sartorius, Göttingen, Germany) to a volume <100 µL. The liposome-protein solution must be resuspended well by pipetting up and down before ultracentrifugation to reduce protein loss through co-pelleting

DPPC and DOPC acceptor liposomes were prepared by 5x ultrafiltration in a 0.2 µm centrifugal concentrator (Vivaspin 500, PES; Sartorius) at 6,000 x g and dilution with liposome buffer. Thorough resuspension between rounds is crucial. This step ensures that acceptor liposomes are at least five-fold smaller than fluorescent donor liposomes and can thus be distinguished in a laser scanning microscope with a 63x/1.4 oil objective.

P116 filled with fluorescent lipids was incubated with acceptor liposomes for 2 h at room temperature with gentle shaking. P116 was then removed by washing four times with liposome buffer via ultrafiltration in a 300,000 MWCO centrifugal concentrator (Vivaspin 500, PES) at 6,000 x g. The process was monitored by measuring the P116 concentration in the flow-through until total protein was recovered and the concentration in the filtrate was zero. A dot blot was used to verify that the concentration of P116 in the filtrate was << 0.5 µg. To account for protein carryover below the sensitivity of a dot blot, a control sample was additionally treated with 75 μg/ml Proteinase K (Promega, Madison, USA) for 1 h at 37°C after ultrafiltration. After the incubation, the proteinase was removed with another round of ultrafiltration in 300,000 MWCO centrifugal concentrator (Vivaspin 500, PES) at 6,000 x g. The resulting sample was subjected to analysis by MALDI-TOF mass spectrometry and laser-scanning microscopy (Table S2). The fluorescence intensity of acceptor liposomes after Proteinase K treatment did not significantly differ (Fig. S4c, Table S2), indicating that ultrafiltration was effective in removing P116.

To account for (i) transfer by unspecific binding of lipids to protein, (ii) spontaneous insertion of fluorescent lipids into the acceptor liposomes from possible donor liposome contaminations of the final sample, or (iii) spontaneous fusion between acceptor and donor liposomes from possible donor liposome contaminations of the final sample, the procedure detailed above was repeated in the presence of P110/140 (ectodomain of a non-lipid binding mycoplasma membrane protein) instead of P116, and the sample was subjected to analysis by laser-scanning microscopy (Table S1). All experiments were carried out in triplicate, and all results could be replicated.

### Laser scanning microscopy

Laser-scanning fluorescence microscopy was performed on an inverse confocal microscope (LSM780, Axio Observer Z.1; Carl Zeiss, Jena, Germany) with a Plan-Aptochromat 63x/1.4 oil objective and Immersol 518F (Carl Zeiss, Jena, Germany) using laser lines at 405 nm and 458 nm. The pinhole was set to 1 airy unit. Imaging was done in an 8 μL drop of sample in an 8-well µ-Slide with glass bottom (ibidi). The fluorescence intensity of individual liposomes was analysed with Zeiss ZEN 2012 (black) v.8.0.5.273 and ImageJ 1.53t (*28*).

### Matrix-assisted laser desorption/ionisation mass spectrometry

All samples were mixed in a 1:1 ratio with sDHB (Super-DHB; Bruker, Billerica, USA) matrix solution (50 mg mL in 50% acetonitrile (ACN), 50% water, and 0.1% trifluoroacetic acid). Subsequently, 1 μL aliquots of the mixture were deposited on a BigAnchor MALDI target (Bruker, Billerica, USA) and allowed to dry and crystallise at ambient conditions. MS spectra were acquired on a rapifleX MALDI-TOF/TOF (Bruker, Billerica, USA) in the mass range of 100–2,000 m/z in reflector positive and negative mode for lipid measurements. The Compass 2.0 (Bruker, Billerica, USA) software suite was used for spectra acquisition and processing.

### Overexpression of native P116 in *M. pneumoniae*

The gene *mpn213* (coding for P116) was amplified from isolated genomic DNA of *M. pneumoniae*strain M129 using primers 5’ CAGGACTCGAGCCGTTTGTTTAAGGACAA-AAC 3’ and 5’ GGAGATATCCTAAAAACCAACGAACCAGAAG 3’, bearing an EcoRV and a XhoI restriction site, respectively. As little is known about Mycoplasma promoters and enhancers, the position of the forward primer was chosen at the end of *mpn212*, upstream of *mpn213*, to include the native promoter and regulatory elements. The *M. pneumoniae* replicating plasmid pGP2756^17^ was linearised by PCR using primers 5’ CTGAGATATCTAGTTATTG-CTCAGCGGTGG 3’ and 5’ CAGGACTCGAGCACTTTTCGGGGAAATGTGC 3’, bearing an EcoRV and a XhoI restriction site, respectively. After restriction with the respective enzymes, the linearised backbone and amplified gene were ligated and used to transform DH5α *Escherichia coli* cells. The resulting plasmid is available upon request.

After verification by sequencing, *M. pneumoniae* cells were transformed as previously described (*29*): *M. pneumoniae* cultures were grown to late-exponential phase in 75 cm^2^ culture flasks. The adherent cell layer was washed three times with a chilled electroporation buffer (8.0 mM HEPES, 272 mM sucrose, pH 7.4), scraped off and resuspended in 0.5 mL of the same buffer. The suspension was passed ten times through a 25-gauge syringe needle. Then, 200 μL aliquots were mixed with 1 μg of the plasmid and transferred to 0.2 cm electro cuvettes (BIO-RAD, Hercules, USA), chilled for 15 min on ice, and electroporated in a BIO-RAD MicroPulser Electroporator using the Ec2 preset (2500 V). After pulsing, the cells were chilled on ice for 15 min before 600 μL of prewarmed SP4 medium was added. Cells were allowed to recover at 37°C for 2 h before the transformation volume was inoculated into a 75 cm^2^ tissue culture flask filled with 20 mL medium containing 10 μg/mL tetracycline and further cultured at 37°C.

### Cryo-electron tomography of *M. pneumoniae*

Adherently growing *M. pneumoniae* cells overexpressing P116 were harvested by scraping 2 days after transformation without washing and pelleting for 20 min at 15,000 x g. The pellet was washed once in 1 mL PBS, re-pelleted and then resuspended in 100 μL PBS. Multiple dilutions of the suspension were prepared and passed multiple times through a 25-gauge syringe needle until all clumps were resolved. The solution was mixed with fiducial markers (Protein A conjugated to 5 nm colloidal gold; Cell Biology Department, University Medical Center Utrecht, The Netherlands) in some cases. From the solutions with or without fiducials, a 3.5 μL drop was applied to a (45 s) glow-discharged R1.2/1.3 C-flat grid (Electron Microscopy Science, Hatfield, USA) with a 4 nm carbon coat, and plunge-frozen in liquid ethane (Vitrobot Mark IV; Thermo Fischer Scientific) at 100% relative humidity, 4°C, and a nominal blot force of –1, with wait and blotting times of 8–12 s.

Tilt-series were recorded using SerialEM v4.0.14 (*30*) at a nominal magnification of 81,000 (1.112 Å per pixel) in nanoprobe EFTEM mode at 300 kV with a Titan Krios G2 (Thermo Fischer Scientific) electron microscope equipped with a BioQuantum-K3 imaging filter (Gatan), operated in zero-loss peak mode with 20 eV energy slit width, and a K3 Summit detector (Gatan). The total dose per tomogram was 120 e^−^/Å^2^, and the tilt series covered an angular range from −60° to 60° with an angular increment of 3° and a defocus set at −3 μm. Local motion correction and CTF estimation of tilt series were done in Warp v.1.0.9 (*31*). Image stack alignment was done in IMOD v.4.11.8 (*32*). Tomograms were reconstructed in Warp v.1.0.9. Particles were clicked on deconvoluted tomograms in ArtiaX v1.0 (ChimeraX extension) (*33*). Sub-tomograms were reconstructed with Warp, using the star files from ArtiaX after adding micrograph names and modifying the header. Initial models from sub-tomograms and 3D refinement were done with Relion v.3.1 (*34*) with 698 homogenous and 1101 heterogeneous particles, a box size of 128 pixels, a pixel size of 4.445 Å/pix, a mask diameter of 450 Å, and a resolution of ∼30 Å (EMD-18629). Filtered and projected sub-tomograms of individual particles were inspected in Amira v.5.3.3 (Thermo Fischer Scientific).

### Cargo binding assay

Previously emptied P116 was incubated with an excess of DPPC liposomes in buffer or 2.8 µL of a 510 µM peptide solution in buffer. The incubation was carried out in 50 µL total volume for 2 h at 37°C with mild shaking. The liposomes were removed by ultracentrifugation for 30 min at 150,000 x g in a Ti70 rotor with 1.5 mL Eppendorf tube adapters and re-ultracentrifugation of the upper 80% of supernatant, before collecting the upper 90% of supernatant and washing it three times in 50,000 MWCO centrifugal concentrators (Vivaspin 500, PES; Sartorius, Göttingen, Germany). Unbound peptide was removed through four rounds of ultrafiltration in a 50,000 MWCO centrifugal concentrator (Vivaspin 500, PES) at 6,000 x g. Efficient removal was verified by MALDI-TOF mass spectrometry of a control repeating the procedure detailed above (i) in the absence of protein (testing for unspecific binding of DPPC or peptide to the filter or tube) (ii) in the presence of P110 and P140, *M. pneumoniae* membrane proteins (testing for nonspecific protein binding effects).

### Single-particle cryo-EM

A 3.5 μL drop of purified P116 (300 μg/mL in 20 mM Tris, pH 7.4 buffer for full-length P116 (30–957) and 3 mg/mL in 20 mM Tris, 0.5 mM CHAPSO, pH 7.4 buffer for ΔN-P116 (246–818)) was applied to a 45 s glow-discharged R1.2/1.3 C-flat grid (Electron Microscopy Science, Hatfield, USA), and plunge-frozen in liquid ethane (Vitrobot Mark IV; Thermo Fischer Scientific) at 100% relative humidity, 4°C, a nominal blot force of –3, a wait time of 0.5 s, and a blotting time of 12 s. Before freezing, Whatman 595 filter papers were incubated for 1 h in the Vitrobot chamber at 100% relative humidity and 4°C.

Dose-fractionated movies of lipid-filled and peptide-filled full-length P116 and lipid-filled ΔN-P116 were collected with SerialEM v4.1.0beta (*30*) at a nominal magnification of 165,000 (0.819Å per pixel) in nanoprobe EFTEM mode at 300 kV with a Titan Krios (Thermo Fischer Scientific) electron microscope equipped with a GIF Quantum S.E. post-column energy filter in zero-loss peak mode and a K2 Summit detector (Gatan, Pleasanton, USA). Micrographs were collected with 50 frames per micrograph and 50 e^−^/Å^2^ exposure. The camera was operated in dose-fractionation counting mode with a dose rate of ∼8 electrons per Å^2^ s^−1^. Details can be found in Table S3a & b, Table S5 and Fig. S11, S12, & S14.

For empty full-length P116 and ΔN-P116 monomers and dimers after emptying and after attempted refilling, dose-fractionated movies were collected using EPU v.3.1.3 (Thermo Fischer Scientific) at a nominal magnification of 105,000 (0.837 Å/pix) in nanoprobe EFTEM mode at 300 kV with a Titan Krios G3i electron microscope (Thermo Scientific) equipped with a BioQuantum-K3 imaging filter (Gatan) and operated in zero-loss peak mode with 30 eV energy slit width. Micrographs were collected with 50 frames per micrograph and 50 e^-^/Å^2^ exposure. The camera was operated in counting mode with a dose rate of ∼15 electrons per Å²s^-1^. Details can be found in Table S3a & b and Fig. S11 & S12.

CryoSPARC (*35*) was used to process the cryo-EM data unless stated otherwise. Beam-induced motion correction and contrast transfer function (CTF) estimation were performed using CryoSPARC’s own implementation. The cryo-EM structure of ΔN-P116 was initially obtained from dimers. However, the dimerisation rate of ΔN-P116 was noticeably less than that of the full-length P116, as evidenced by the presence of approximately 50% monomers in the 2D classes of purified ΔN-P116 and >90% monomers in the 2D classes of ΔN-P116 after emptying, compared with only dimers in the 2D class averages of the full-length P116 after purification and ∼30% monomers in the 2D classes of full-length P116 after emptying. We consider it probable that the observed decay of the dimer is an artefact of the emptying with Triton X-100, through which the protein is forced to remain structurally destabilised for a prolonged time. When *empty*, P116 is generally conformationally unstable when compared to the *full* state (*15*). To improve the resolution of the *empty* ΔN-P116, we also determined the structure from monomer particles, as they have less conformational freedom than dimers due to their intermonomer flexibility. Importantly, despite having a lower resolution, the density map obtained from the dimers in the dataset matched the conformation of the monomers (EMD-51410, Fig. S9b & S12c, Table S3b & S3c).

For the dimers, particles were initially clicked with the Blob picker using a particle diameter of 200–300Å and extracted with a box size of 512 pixels, binned to 256 pixels. For the monomers, particles were initially clicked with the Blob picker using a particle diameter of 100–200 Å and extracted with a box size of 256 pixels. Particles were then subjected to unsupervised 2D classification. For the final processing, the generated 2D averages were taken as templates for the automated particle picking. We removed false-positive picks by two rounds of unsupervised 2D classification. The remaining particles after 2D classification were used to generate an *ab initio* reconstruction with three classes followed by a heterogeneous refinement with three classes. For the remaining particles after 3D classification, the beam-induced specimen movement was corrected locally. The CTF was refined per group on the fly within the non-uniform refinement^22^. As initial volumes for the non-uniform refinement of the monomers, PDB: 8A9A (filled P116) and PDB:8A9B (empty P116) were truncated to residues 246–818 and filtered to 30 Å. Both maps yielded similar refined structures. Details can be found in Table S3a & b, Table S5 and Fig. S10, S12, & S15.

### Determination of P116 conformation/filling state

In the full state, the cavity typically contains densities from lipid molecules. In the absence of lipid cargo, helix pair 2 (aa 444 - 476) moves towards the inside of the cavity, and the volume of the hydrophobic cavity is reduced by 60% when compared to the *full* state (*15*) (Fig. S1c). To determine the conformation of P116 objectively, the cryo-EM density maps at a defined contour level were rigid-body fitted (chimerax *fitmap*) with both PDB: 8A9B (*empty* conformation of P116) and PDB: 8A9A (*full* conformation of P116). Subsequently, the correlation of the map-to-map fit of helix pair 2 (aa 444 – 476) at 3 Å resolution into the density map was measured (chimerax *molmap* and chimerax *measure correlation*). All correlation values can be found in Table S4.

### Molecular dynamics simulations

In each of the coarse-grained (CG) simulations we used the following workflow except otherwise stated. We solvated each system using the *insane.py* script (*36*) with a ion concentration of 0.15 M NaCl. We ran a 5000-step soft-core (*free-energy = yes*) minimisation followed by a 3000-step hard-core (*free-energy = no*) minimisation, both using the steepest descent algorithm.

For membrane systems, we ran six rounds of NPT equilibration per system at a reference temperature of 310 K (v-rescale thermostat, τ = 1 ps (*37*)) and a reference pressure of 1 bar (Berendsen barostat, semi-isotropic coupling type, τ = 5 ps (*38*)) with ever-decreasing force constant on the position restraints of the membrane. When the configurations used to initialized the simulations were taken froma previous production we ran only the one NPT equilibration step with no position restraints on the membrane.

For the systems in solution, we ran an NPT equilibration at a reference temperature of 310 K (V-rescale thermostat, τ = 1 ps) and a reference pressure of 1 bar (Berendsen barostat, semi-isotropic coupling type, τ = 5 ps).

We then simulated the systems in NPT ensemble using GROMACS 2021.4 (*39*, *40*) at a reference temperature of 310 K (v-rescale thermostat, τ = 1 ps) and a reference pressure of 1 bar (Parrinello-Rahman barostat, semi-isotropic coupling type, τ = 5 ps (*41*)). We used a time step of 20 fs. Coulombic interactions were treated with PME and a cut-off of 1.1 nm as for van der Waals interactions.

In the following paragraphs we discuss modelling choices and further details for each system.

### CG *empty* P116 dimer on membrane

First, we generated the MARTINI CG P116 dimer on a symmetrical bilayer membrane (30% DOPC, 30% DOPE, 40% DOPG) using the Charmm-GUI MARTINI maker webserver (bilayer builder) with the MARTINI 3.0 forcefield (*42–44*) on the PDB structure 8A9A (UniProt serial numbering, residues 60–868 per chain). We prepared two different models. One retained the *interdomain* elastic network (EN) generated by the software (via *martinize2*) to keep the N-terminal domain fixed as seen in the cryo-EM structures (PDB 8A9A, 8A9B) (model **Restrained empty on membrane** in Table 1). For the second model (model **Empty on membrane** in Table 1), we modified the EN to retain only *intradomain* bonds (we later used this criterion for all the CG simulations except otherwise stated). Additionally, we prepared another model where we included the C-terminal stretch of about 100 amino acids to obtain the P116 60-957 model as used in the biophysical assays (model **Empty with C-terminus** in Table 1). To do so, in the absence of an experimental structure, we used the AlphaFold Colab (*45*) prediction of the C-terminal stretch (UniProt serial numbering 868-957 per chain). We merged the P116 60-868 model to two copies of the C-terminus predicted structure using UCSF Chimera (*46*). We modified the EN of this model to retain only the *intradomain* bonds.

**Table 1:** Models used for MD simulations, simulated time and number of runs.

| MODEL | SIMULATED TIME<br>[μs] | N°<br>RUNS |
| --- | --- | --- |
| Full with Cholesterol | 0.6 | 1 |
| Empty on membrane | 33 | 1 |
| Restrained empty on membrane | 21 | 1 |
| Empty with pulled lipids | 35 | 1 |
| Empty in solution | 22 | 1 |
| Restarined DCA | 10 | 1 |
| Restrained empty in solution | 4 | 1 |
| Empty docked with splayed lipid | 14 | 49 |
| Full on membrane | 36 | 1 |
| Empty with C-terminus | 14 | 4 |
| Less full on membrane | up to 53 | 4 |

To incentivise protein-membrane interactions, we set a box height of 17 nm. This choice enables P116 to roam freely in about 13 nm (height) of solvent, which mimics the distance from the membrane allowed by the tethering to *Mycoplasma* as estimated through cryo-ET (see Fig. S7b). We kept this choice consistent for all systems with a membrane.

To avoid unphysical behavior during production of the equilibrated P116 60-957, we first enabled the hydrophobic cavity to collapse while keeping the 868-957 loop restrained in place.

The comparison of the simulated dynamical ensembles with the tomograms of *M. pneumoniae* revealed the incompatibility of the P116 60-868 with *interdomain* EN and P116 60-957 ensembles (see Fig. S6c & d), while the P116 60-868 without interdomain EN showed remarkable agreement (see Fig. 4b and Fig. S6c &d). Thus, we chose the P116 60-868 model without interdomain EN for the MD simulations as it is predictive of the P116 dynamics.

The simulation of P116 60-868 showed docking of the protein to the membrane. We took a docked configuration obtained with the previous run and used it to reinitialise the system where three lipids in proximity of the DCA of the docked P116 monomer were pulled towards the DCA using UCSF Chimera (model **Empty with pulled lipids** in Table 1). This was the first attempt to favor the lipid uptake. We simulated this model for about 35 µs. During this simulation, P116 steadily docked to the membrane via residues F214 and F227 (see Fig. 5a).

### CG *full* P116 dimer on a membrane

Using the same procedure used for the *empty* P116, we obtained the CG model of PDB 8A9A (UniProt serial numbering, residues 60–868 per chain). We proceeded to fill the hydrophobic cavity with lipids (2 DOPCs, 1 DOPEs, 6 DOPGs per cavity) using UCSF Chimera (model **Full on membrane** in Table 1). We maintained the parameters and equilibration protocol consistent but used GROMACS 2022.4.

### CG *less full* P116 dimer on membrane

We extracted a 30°-wringing configuration from the trajectory of the model Full on membrane and removed 3 lipids per cavity from this model. We obtained the model “*Less full on membrane”* (see Table 1). We solvated the model using the *insane.py* script. We maintained the parameters and equilibration protocol consistent but used GROMACS 2022.4. We ran 4 replicas of the system (R1-4). R1 ran for 53 µs, R2 and R3 for 37 µs, and R4 for about 20 µs.

### CG *empty* P116 dimer in solution with free DOPG, DOPE and DOPC lipids

Using the topology of the P116 dimer and lipids from the simulations with membrane and the equilibrated P116 dimer configuration, we created the model of the P116 dimer with sealed hydrophobic cavity surrounded by 15 DOPGs, 7 DOPCs and 7 DOPEs using UCSF Chimera. We prepared two models: one with *interdomain* EL (model **Restrained empty in solution** in Table 1) and one retaining only *intradomain* EL bonds (model **Empty in solution** in Table 1). We then repeated the preparation workflow, but we introduced three elastic bonds (500 kJ/mol) at the opening of the DCA in the **Empty in solution** model (model **Restrained DCA**). The bonds were defined as follows (ID bead 1, ID bead 2, equilibrium distance [nm]):

1. 740, 833, 0.48394
2. 744, 829, 0.48280
3. 742, 831, 0.76426.

### Atomistic simulation of the P116 dimer filled with cholesterol

We created an atomistic model of the P116 dimer (UniProt serial numbering, residues 60– 868 per monomer) with each hydrophobic cavity filled with 12 cholesterols using UCSF Chimera (model id **Full with Cholesterol** in Table 1). We solvated the system with TIP3P water using GROMACS. We minimised the system using the steepest-descent algorithm for 5000 steps and performed a 200-ps-long NVT equilibration using the V-rescale thermostat with a reference temperature of 310 K (τ = 0.1 ps). We simulated the system in the NPT ensemble for 0.6 µs using the Charmm36m forcefield (*47*, *48*), a reference temperature of 310 K (τ = 1 ps, V-rescale thermostat), a reference pressure of 1 bar (τ = 5 ps, Parrinello-Rahman barostat) and a NaCl concentration of 0.15 M. For both the van der Waals (Verlet) and the Coulomb forces (Particle Mesh Ewald), we used a cut-off of 1.1 nm. We chose a 2 fs time step. We used GROMACS 2021.4.

### CG *empty* P116 dimer on membrane with one splayed lipid tail

Based on a frame from the simulation of the *empty* P116 dimer steadily N-docked on the membrane, we created a model where one lipid tail was splayed to be in between the DCA helices and the other tail was inserted in the membrane bilayer. One lipid was deleted from the previous model to allow the positioning of the splayed lipid using UCSF Chimera (model **Empty docked with splayed lipid** in Table 1). We ran 49 replicas of this system, which resulted in 41 replicas extracting the lipid and 8 where the lipid did not move or escaped. We computed the statistics based on a 1 nm distance threshold between the lipid and the cavity inner dorsal side.

### Definition of arc diameter, wringing angle, interdomain angle, accessible entry size and reference structures

We defined the *arc diameter* of P116 as the average of the distances between residues 416 of chain A and 253 of chain B and between residues 253 of chain A and 416 of chain B per frame.

We defined the *wringing angle* as the angle between the planes containing the centres of mass of residues 280 and 301 and of the whole core domain of each monomer. We defined as offset the wringing angle achieved when the two monomers were both facing approximately downwards (in general, in the same direction). We then subtracted this value from the raw values to set the *up-up conformation* to 0°. Accordingly, we find that the *up-down conformation* is at 95°.

We defined the *interdomain angle* as the angle between the segments connecting the centers of mass of

- the residue pairs 164 and 181, and 107 and 108 and
- the residue pair 382 and 404 and the group 708 to 712.

We defined the entry size by selecting 3 pairs of backbone beads (BB) per entry:

- F1: the BB of residue pairs (471, 519), (470, 520), (469, 521).
- F2: (458, 467), (455, 469), (451, 473).
- F3: (518, 541), (522, 536), (525, 533).
- DCA: (374, 417), (376, 415), (375, 416).

We then computed the average distance per frame and subtracted the bead size to account for the Lennard-Jones radius (0.47 nm).

We defined the MARTINI CG models of the cryo-EM structures (*full* P116 with PDB 8A9A, *empty* P116 with PDB 8A9B, and *refilled* P116 EMD-15276) as reference structures. All reported values are the mean (for the wringing angle and the arc) or the median (for the interdomain angle).

### Statistical Analysis

The lipid transfer assay was performed in three biological replicates with similar results. Fluorescence microscopy images were recorded and analyzed by to individual investigators, one of them blinded. Differences of fluorescence intensities between the samples were analyzed by unpaired Student’s *t* test.

All mass spectrometry results are representative of two biological replicates with similar results. The investigator analyzing the mass spectrometry data was blinded. For cryo-EM data, all particles were included. Investigators were not blinded, as data analysis was performed computationally with cryoSPARC. Unsupervised 2D classification was used to cluster the data sets. Refinement and averaging were done with random individual half sets. All experimental findings could be replicated.

Except otherwise stated, all reported values derived from MD trajectories are reported as the mean ± standard deviation (STD, computed assuming a gaussian distribution) (for the wringing angle and the arc diameter) or the median ± STD (for the interdomain angle).

## Acknowledgments

We thank the Frankfurt Center for Electron Microscopy and the Frankfurt Center for Advanced Light Microscopy for measurement time. We thank the Central Electron Microscopy Facility at the MPI of Biophysics in Frankfurt, which enabled us to collect the tomography dataset as well as the cryo-EM dataset of *empty* P116 monomers, particularly S. Welsch and S. Prinz who assisted during the data collection. We thank N. Morgner and J. Schulte for their support during LILBID mass spectroscopy measurements, C. Glaubitz and I. Weber for assistance with liposome preparation, I. Fita, D. Vizarraga and J. Martín for the kind provision of purified P116 (246–818), M. Kunz for support with tomogram reconstruction, and A. N. Birtasu and U. H. Ermel for assistance with sub-tomogram averaging. We thank Jörg Stülke *for providing the M. pneumoniae* replicating plasmid pGP275617, I. Fita/J. Martín for gifting us the vector pOPINE_P116, and S. Knapp/M. Schwalm for providing a hydrophobic peptide. We thank M. R. Vabulas for valuable discussions and comments.

## Funding

Goethe University Frankfurt (SMA, RC)

Frankfurt Institute of Advanced Studies (SMA, RC)

LOEWE Center for Multiscale Modelling in Life Sciences state of Hesse (SMA, RC)

German Research Foundation, CRC 1507: Membrane-associated Protein Assemblies, Machineries, and Supercomplexes (P09) (Project ID 450648163) (SMA, RC)

International Max Planck Research School on Cellular Biophysics (SMA, RC)

Center for Scientific Computing of the Goethe University (SMA, RC)

Jülich Supercomputing Centre (SMA, RC)

German Research Foundation, grant FR 1653/6-3 (ASF, LS)

German Research Foundation, SFB 902 (ASF, KW, DB)

German Research Foundation, Research Training Group iMOL, grant GRK 2566/1 (SM)

## Author contributions

Conceptualization: SM, SMA

Data curation: SM, SMA, DB

Formal analysis: SM, SMA

Funding acquisition: RC, ASF

Investigation: SM, SMA, LS, JMC, KW, DB

Supervision: RC, ASF, JL

Visualization: SM, SMA

Writing – original draft: SM, SMA

Writing – review & editing, SM, SMA, RC, ASF

## Data and materials availability

The cryo-EM structures determined in this study are available at http://www.emdataresource.org/ under EMD-50314, EMD-51408, EMD-51409, EMD-51412, EMD-51411, EMD-51410 and EMD-18476 (Table S3a & b, Table S5). The MD data supporting this study’s findings, including Videos S1 and S2, are available in MD_P116_22042025.zip at https://doi.org/10.5281/zenodo.13383577.

