## Supplemental Information for "How does *Mycoplasma pneumoniae* steal lipids from its host membranes?"

**This PDF file includes:**

Figs. S1 to S14
Tables S1 to S5
Movies S1 to S2

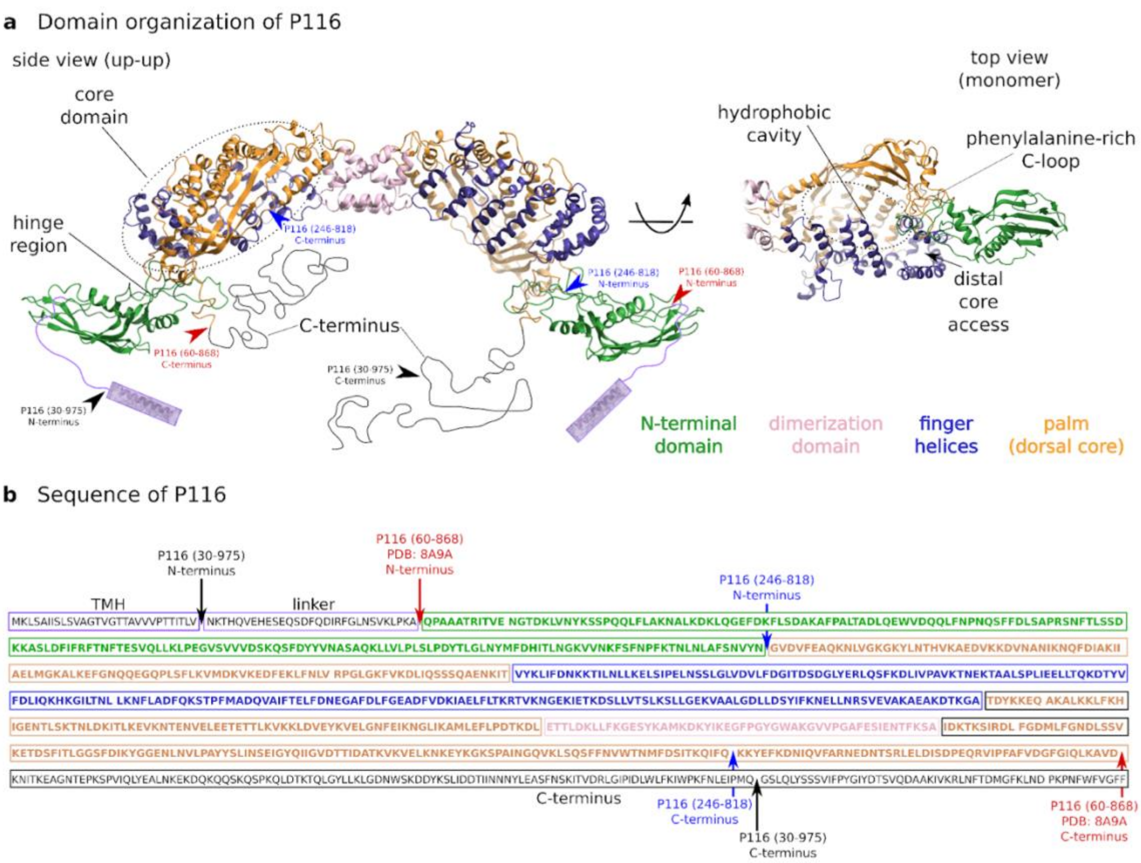

**Fig. S1. Domain organisation and sequence of P116 fragments.** (**A**) Ribbon model of P116 (PDB: 8A9A). The domains are color-coded (N-terminal domain in green, core domain in blue (for the four amphipathic helices) and orange (for the palm), dimerisation domain in pink, transmembrane helix in purple, linker in light purple and C-terminus in black) and the ectodomain fragments used in this study are indicated with black (P116 30-975), red (P116 60-868) and blue (P116 246-818) arrows. The transmembrane helix, linker and C-terminus are not structurally resolved and therefore only sketched into the model. (**B**) Amino acid sequence of P116 with the same color scheme that was used for the ribbon model. The ectodomain fragments used in this study are indicated with black (P116 30-975), red (P116 60-868) and blue (P116 246-818) bars.

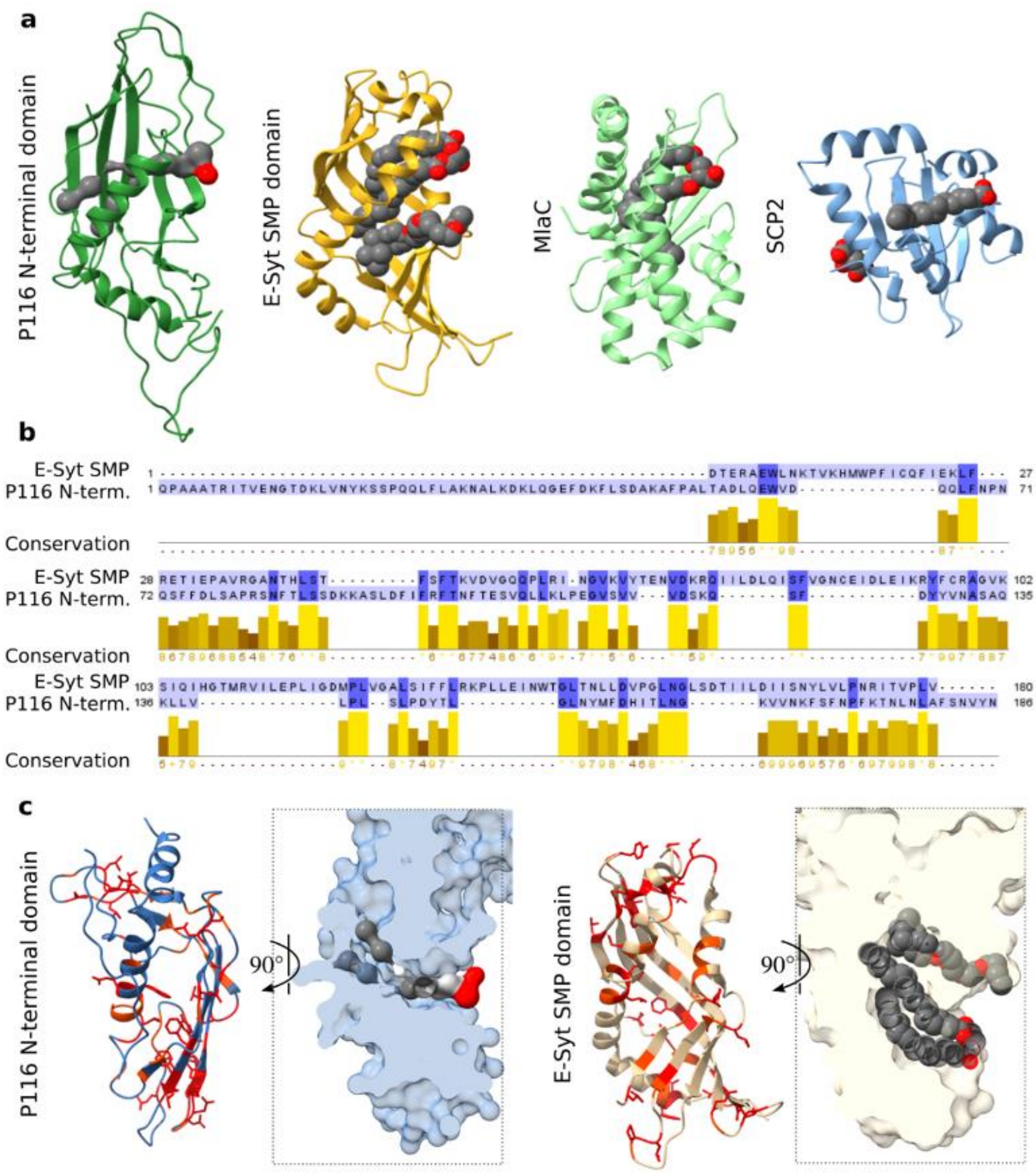

**Fig. S2. Structural comparison of P116 N-terminal domain with small lipid shuttle proteins.** (**A**) The folds of the P116 N-terminal domain (PDB: 8a9a), the E-Syt SMP domain (PDB: 4P42), MlaC (PDB: 5UWA) and SCP2 (PDB: 4JGX) share similarities: an antiparallel β-sheet covers three sides of a hydrophobic channel, while the remaining side is closed by one or more α-helices. Proteins are shown as ribbon models, and lipids are shown as grey sphere models. (**B**) Sequence alignment of the P116 N-terminal domain and the E-Syt SMP domain. The sequences are colored by percentage identity. The alignment has 14% sequence identity, 25% similarity and 55% gaps. (**C**) Detailed structural comparison of the P116 N-terminal domain and the E-Syt SMP domain. Identical amino acids are shown in red as stick models; closely related amino acids are shown in orange. Lipid tails are colored grey, and lipid head groups are colored red.

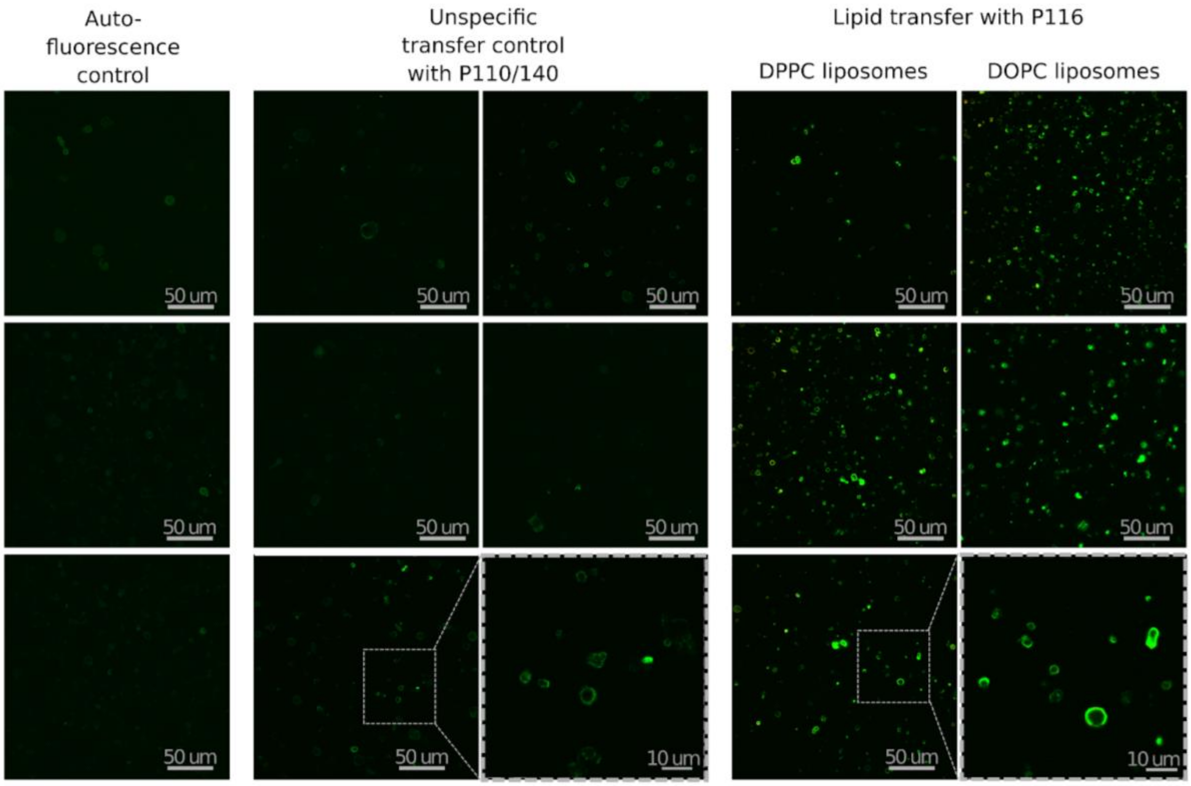

**Fig. S3. Confocal light microscopy images of delivery into DPPC liposomes.** Merge of Dansyl and NBD channels. Left column: Acceptor liposomes (auto-fluorescence control). Middle column: Workflow without P116 to control for spontaneous fusion/transfer control. Right column: DPPC and DOPC Vesicles incubated with P116, according to the workflow shown in Fig. 1. The experiment was done in triplicate. A statistical analysis of the liposome fluorescence can be found in Fig.1 and Table S2.

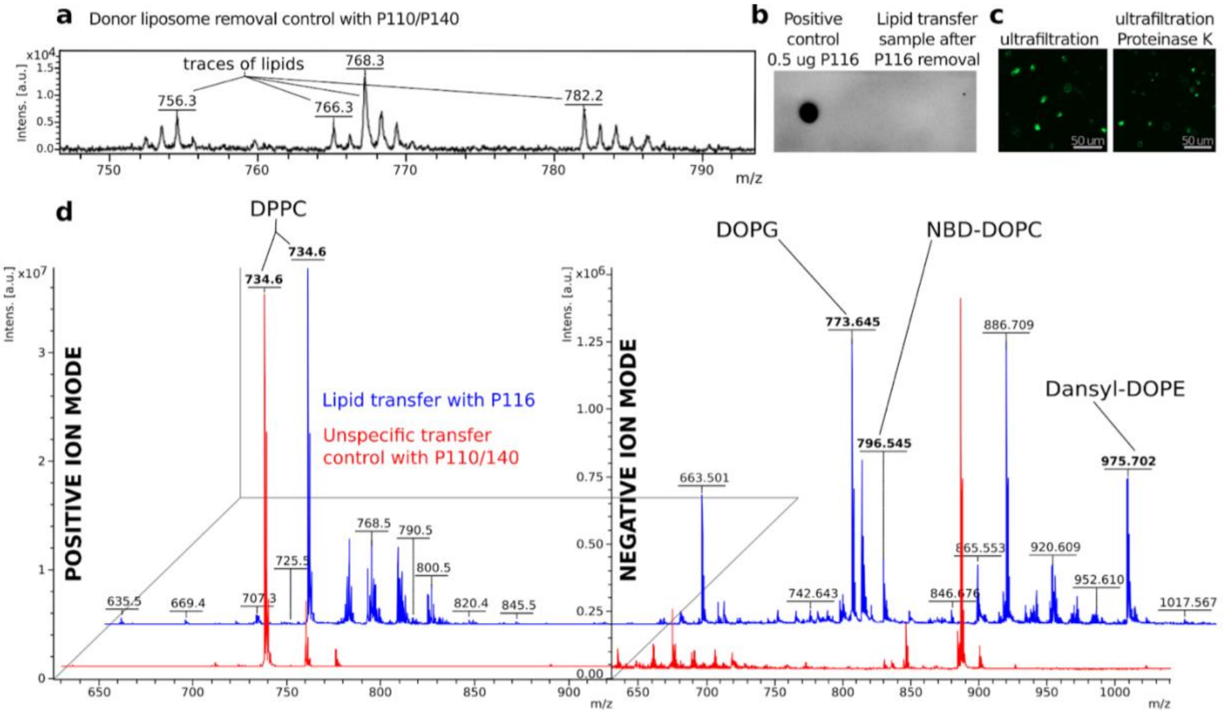

**Fig. S4. Controls for the lipid transfer assay.** (**A**) Mass spectra of the supernatant after two rounds of ultracentrifugation of a donor liposome solution mixed with the non-lipid binding mycoplasma proteins P110/P140. Traces of lipids at the masses 756, 766, 768 and 782 Da are visible. DOPG, NBD-DOPC and Dansyl-DOPE are absent. While the removal method seems adequate, carryover under the detection limit cannot be ruled out. Thus, we included an additional control for the spontaneous fusion of carryover donor liposomes. (**B**) Dot blot of filtrate after four rounds of ultrafiltration of DPPC acceptor liposomes mixed with P116. The dot blot demonstrates that the potential carryover of fluorescently filled P116 into the final sample was far below 0.5 µg. (**C**) Confocal light microscopy images of the lipid transfer sample after removal of P116 with ultrafiltration (left) and after removal of P116 with ultrafiltration and an additional Proteinase K digest. The difference between the average fluorescence intensity was not significant (see Supplemental Table 4), indicating that removal with ultrafiltration only is sufficient. (**D**) Analysis of DPPC acceptor liposomes after treatment according to the workflow detailed in Fig. 1a (lipid transfer with P116, blue) and after treatment according to the same workflow with non-lipid binding mycoplasma proteins P110/140 instead of P116 (unspecific transfer control, red) using mass spectrometry. In positive ion mode (right), a clear peak at 734 Da, the mass of DPPC, is visible in both samples. In negative ion mode, lipids at the masses 773, 796 and 975 Da are visible, corresponding to DOPG, NBD-DOPC and Dansyl-DOPE, respectively. Masses of other lipids (e.g. at 663 and 886 Da) are likewise visible. Those masses potentially originate from lipids that were still present in P116 after the emptying procedure or from contamination of the sample.

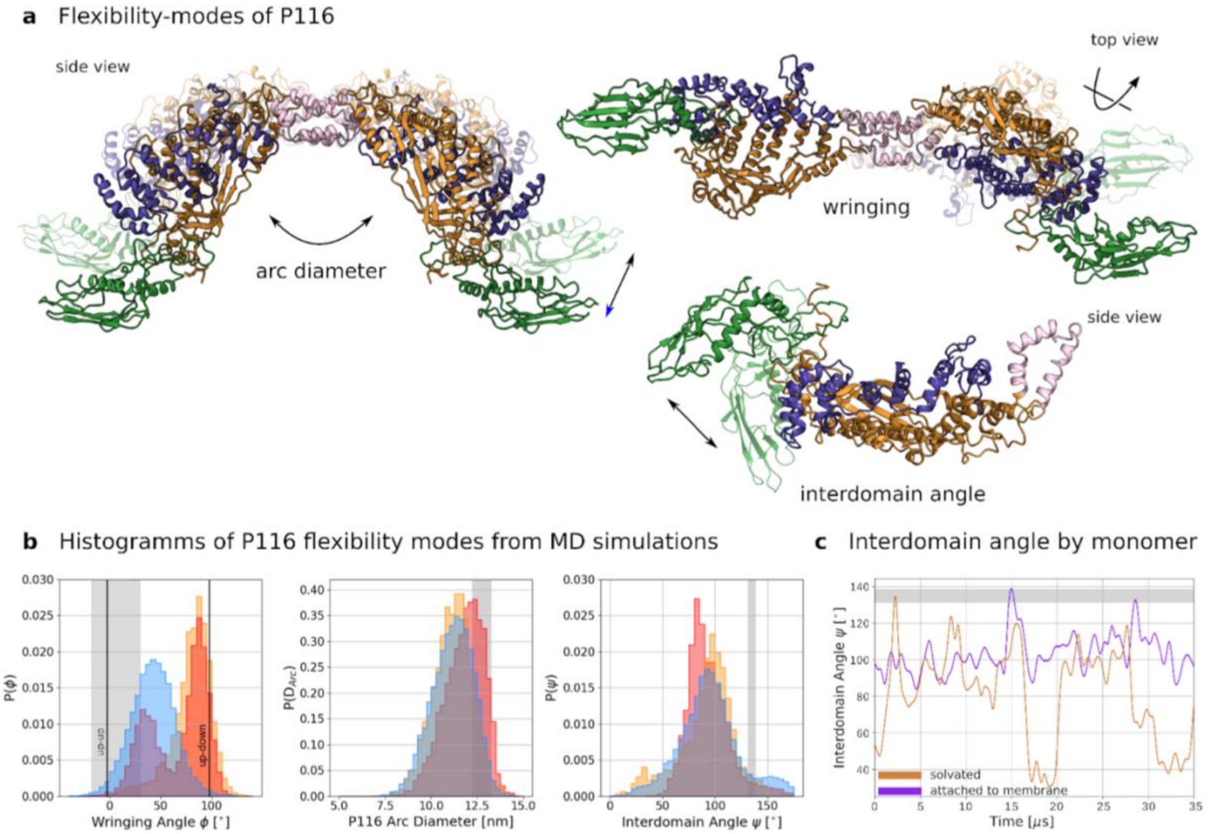

**Fig. S5. Conformational flexibility of P116 in cryo-EM and MD simulations.** (**A**) Flexibility-modes of P116 from single particle cryo-EM of the ectodomain (P116 60-868, PDB: 8a9a and 8a9b) in solution. The arc diameter (distance between monomers) ranges between 11.5 and 12.5 nm. The rotational angle between the monomers (wringing angle) ranges between 0° and 45° (measured on one monomer with the second monomer fixed). The incline angle between the N-terminal domain and the core domain (interdomain angle) remained between 120° and 135°. The definition and calculation method of arc diameter, wringing angle, and interdomain angle can be found in the Material and Methods section. (**B**) Histograms of P116 flexibility modes from MD simulations, including *empty* P116 in solution (blue histograms) and on a membrane (orange histograms) and *full* P116 on a membrane (red histograms). Overall, the histograms indicate greater flexibility than the range of motion observed in cryo-EM structures of solvated P116 (indicated by grey bars). The cargo-dependent population at a wringing angle of ~30° corresponds with the wringing angle observed in the cryo-EM structures of *refilled* P116 (EMD-15276, dashed line). ***Left*** Wringing angle: Simulated *empty* P116 in solution displayed an average wringing angle of 40° ∓ 20°, allowing the monomers to face in the same direction (up-up). In contrast, *empty* P116 on a membrane displayed a far increased wringing angle of 80° ∓ 20°, which enables the monomers to face opposite directions (up-down) (Figure 3ba, *left*). Simulated *full* P116 on a membrane displayed a similar population around ~80° (90° ∓ 10°) but also a second population around 30° ∓ 10°, which corresponds to the wringing angle observed in the *refilled* cryo-EM structure (EMD-15276). ***Middle*** Arc diameter: simulated *empty* P116 in solution and on a membrane displayed an average arc diameter of 11 ∓ 1 nm, which fits the 11.5 nm arc diameter of the *empty* P116 cryo-EM structure (PDB: 8A9B). Simulated *full* P116 on a membrane displayed a slightly shifted arc diameter population around 12 ∓ 1 nm, fitting the 12.5 nm arc diameter of the *full* P116 cryo-EM structure (PDB: 8A9A). ***Right*** Interdomain angle: simulated *empty* P116 in solution and on a membrane displayed an interdomain angle around 90° ∓ 30° and 100° ∓ 30°, respectively. Simulated *full* P116 on a membrane displayed a sharper peak around 90° ∓ 20°.  In general, the observed ranges were much broader than those of the cryo-EM structures, where the interdomain angle lies between 120° and 135°. (**C**) Interdomain angle (time series): the angle is different between the monomers when one is stably bound to the membrane, i.e., the monomer attached to the membrane (purple) has a fixed configuration of the N-terminal domain. In contrast, the N-terminal domain of the solvated monomer (orange) is free to move.

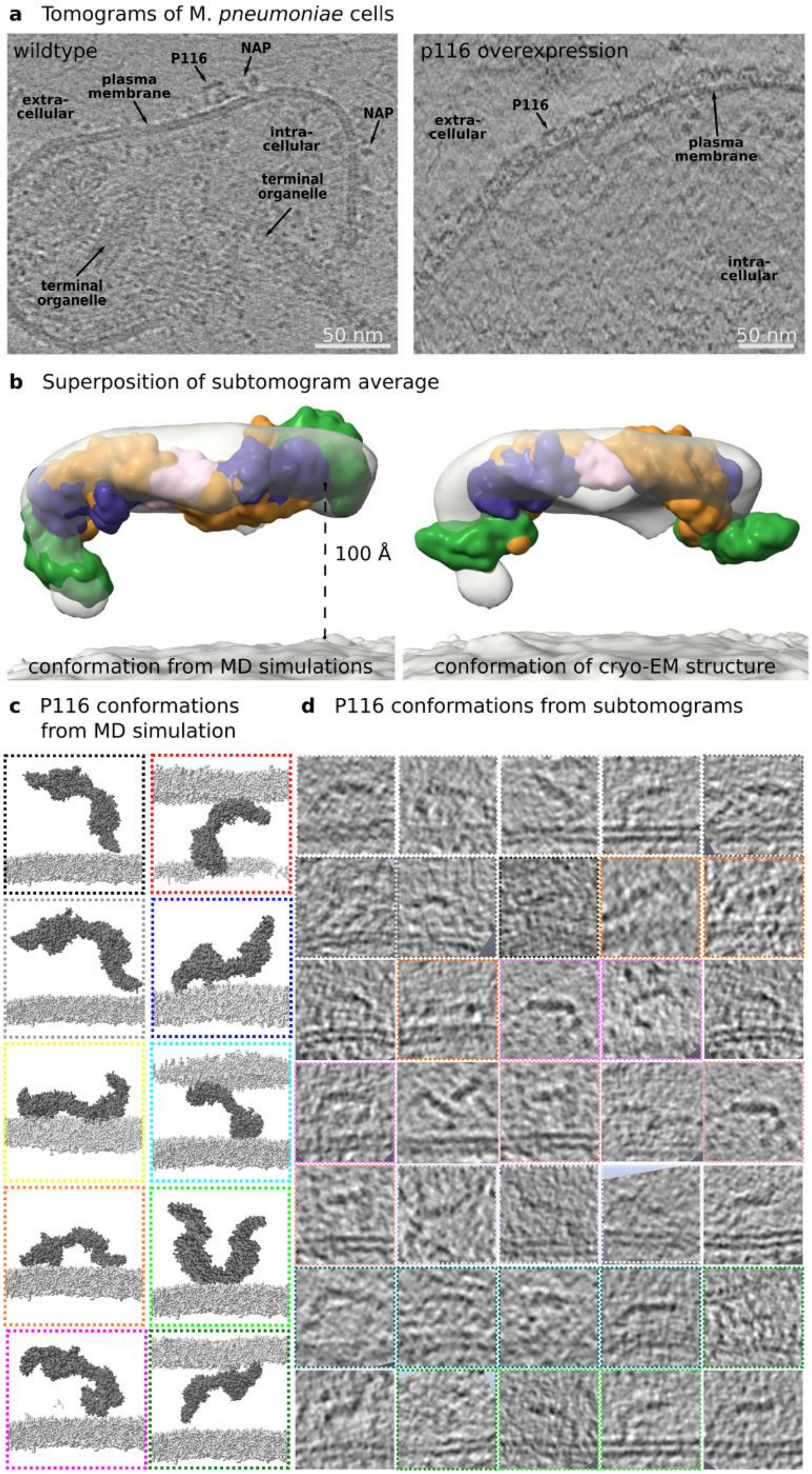

a Tomograms of M. pneumoniae cells
wildtype
P116
NAP
plasma membrane
extra-cellular
intra-cellular
terminal organelle
terminal organelle
50 nm
p116 overexpression
P116
extra-cellular
plasma membrane
intra-cellular
50 nm
b Superposition of subtomogram average
100 Å
conformation from MD simulations
conformation of cryo-EM structure
c P116 conformations from MD simulation
d P116 conformations from subtomograms

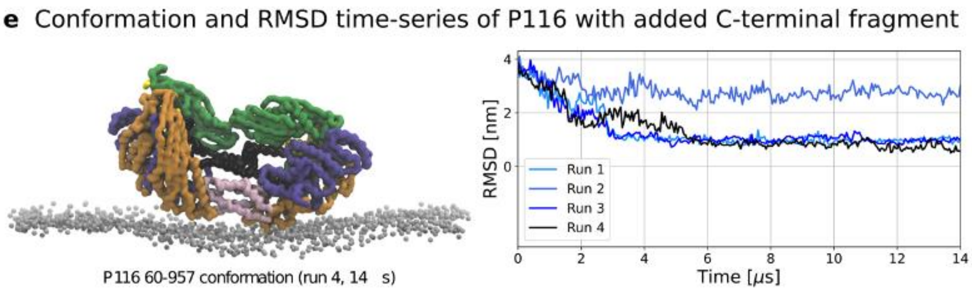

**Fig. S6. Cryo-ET analysis of P116 anchored on the mycoplasma membrane.** (**A**) Computational slices through tomographic reconstructions (0.837 Å/pix) of (left) a wildtype *M. pneumoniae* cell and (right) a cell overexpressing *p116* from its native promoter. In the computational slices, the characteristic features of *M. pneumoniae* cells can be seen, such as the terminal organelle in the cytoplasm and several adhesion complexes (NAP) at the plasma membrane. In the wild-type cell, just one individual P116 particle can be discerned. In comparison, the cell overexpressing *p116* is tightly decorated with P116 particles.
(**B**) 30 Å average of 700 particles (grey surface, EMD-18629). The particle set used for the average contained conformationally homogenous side views. The overall tomographic particle set was too conformationally heterogeneous to average, and particle numbers were not sufficient for classification of conformations beyond the averaged conformation shown. On the left, superposition of the sub-tomogram average with P116 in a conformation obtained from MD simulations. This conformation has a pronounced wringing angle between the monomers and a steep incline angle between the core and N-terminal domains. On the right, superposition of the sub-tomogram average with the single-particle cryo-EM structure in solution (PDB: 8a9a and 8a9b). The single-particle structure has a limited range of motion that does not allow pronounced wringing and high incline angles. It does not fit the sub-tomogram average well. (**C**) Conformations of P116 (60-868) derived from coarse-grain MD simulations.
(**D**) Sub-tomographic projection slices of individual particles in various conformations on the membrane. The colored outlines indicate similarities with simulated conformations from panel c.
(**E**) On the left, the render shows a representative frame corresponding to the conformation explored by P116 (60-957) on the membrane in 3 out of 4 runs of MD simulation. The C-terminal loop, shown in black, preferably inserts into the hydrophobic cavity. On the right, the RMSD time series of the four simulated trajectories computed using the simulation frame shown in the render as a reference configuration.

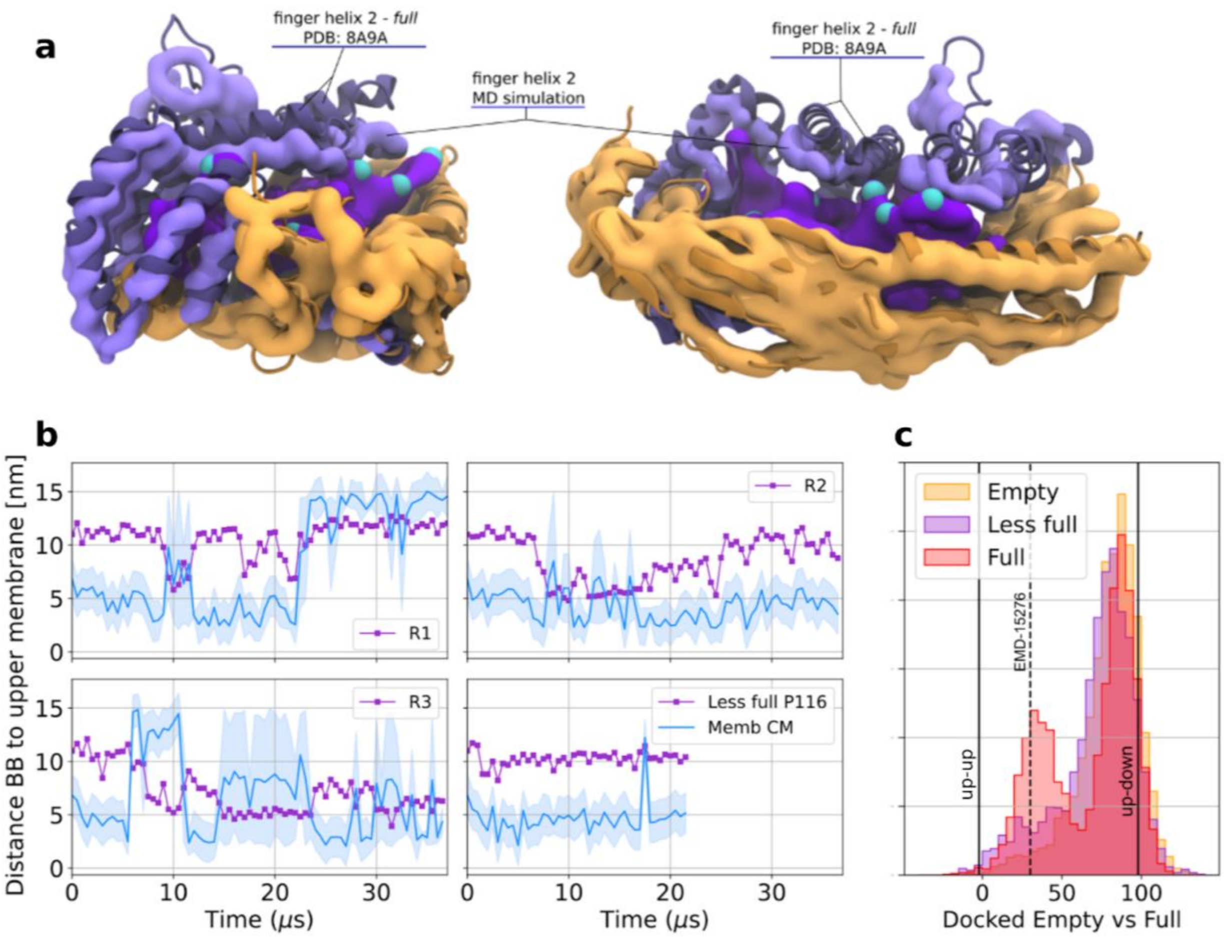

**Fig. S7. P116 undocking mechanism is regulated by the filling state.** (**A**) Cargo mass used in MD simulations: Superimposition of an MD simulation render of the full P116 model with a ribbon model of the full cryo-EM structure (8A9A and EMD-15276). The model used for MD simulation contains less cargo. Lipids are shown as purple and blue spheres. (**B**) Time series of each replica (R1-4) of the less full P116 model on a membrane. The purple square markers and line show the center of mass position of the phenylalanine-rich loop (aa848-868) along the box height. The solid blue line shows the instantaneous position of the center of mass of the membrane along the box height. The shaded blue area delineates the upper and lower leaflets' positions in the box. Thus, whenever the shaded area widens abruptly, it signifies that the membrane bilayer is split between the top and bottom of the box. Therefore, the crossing of the purple line with the shaded area does not correspond with a docking event. The only docking event is visible after about 20 μs of R1, where the purple line is consistently close to the membrane's lower leaflet. (**C**) Distribution of the wringing angle for empty, less full and full P116 simulated models. The population around 30° increases with the increasing mass of the cargo. The sample trajectories used for the comparison are about 30 μs long each. To correctly compare the populations among the models, we extended the R1 trajectory of the less full model to obtain at least 30 μs during which P116 was membrane-bound.

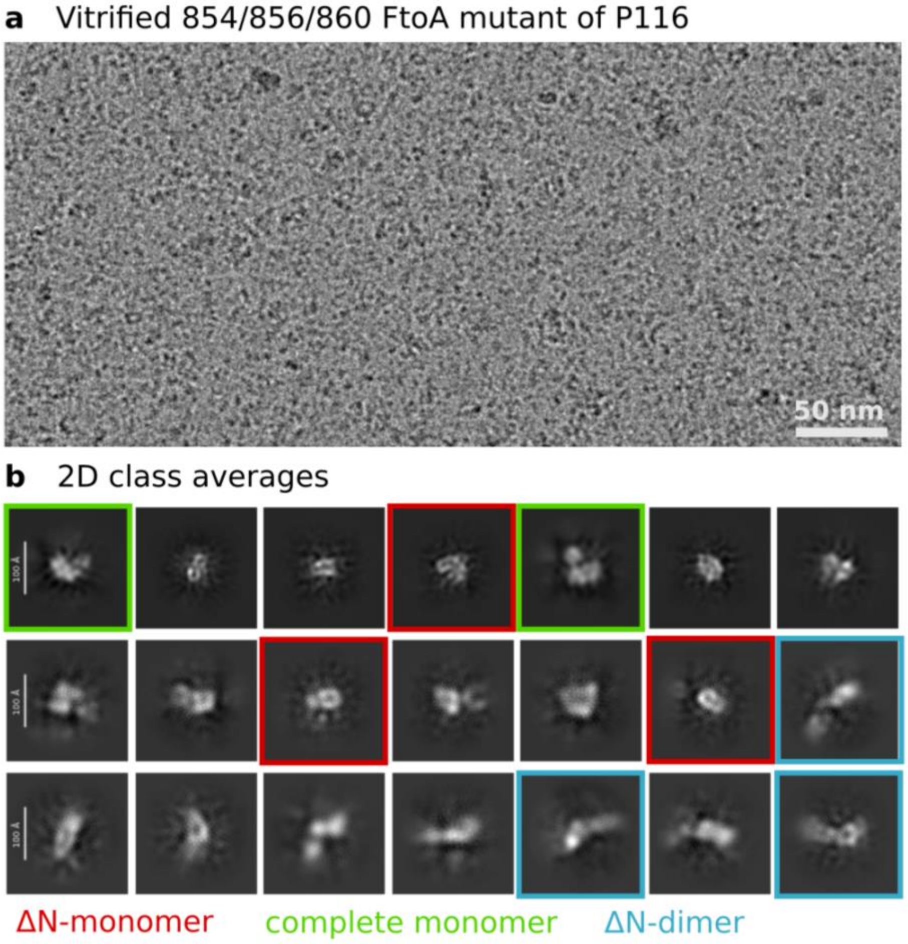

**Fig. S8. Cryo-EM data acquisition and processing of 854/856/860 FtoA mutant of P116.** (**A**) Cryo-electron micrograph of the vitrified mutant in the presence of 0.1% LMNG. In contrast to the wild-type ectodomain of P116, the mutant required the use of detergent to prevent aggregation. (**B**) Exemplary 2D class averages of the mutant showing high structural heterogeneity resulting in low-resolution averages. We observed both monomer and dimer classes, of which many appeared to be missing the entire N-terminal domain.

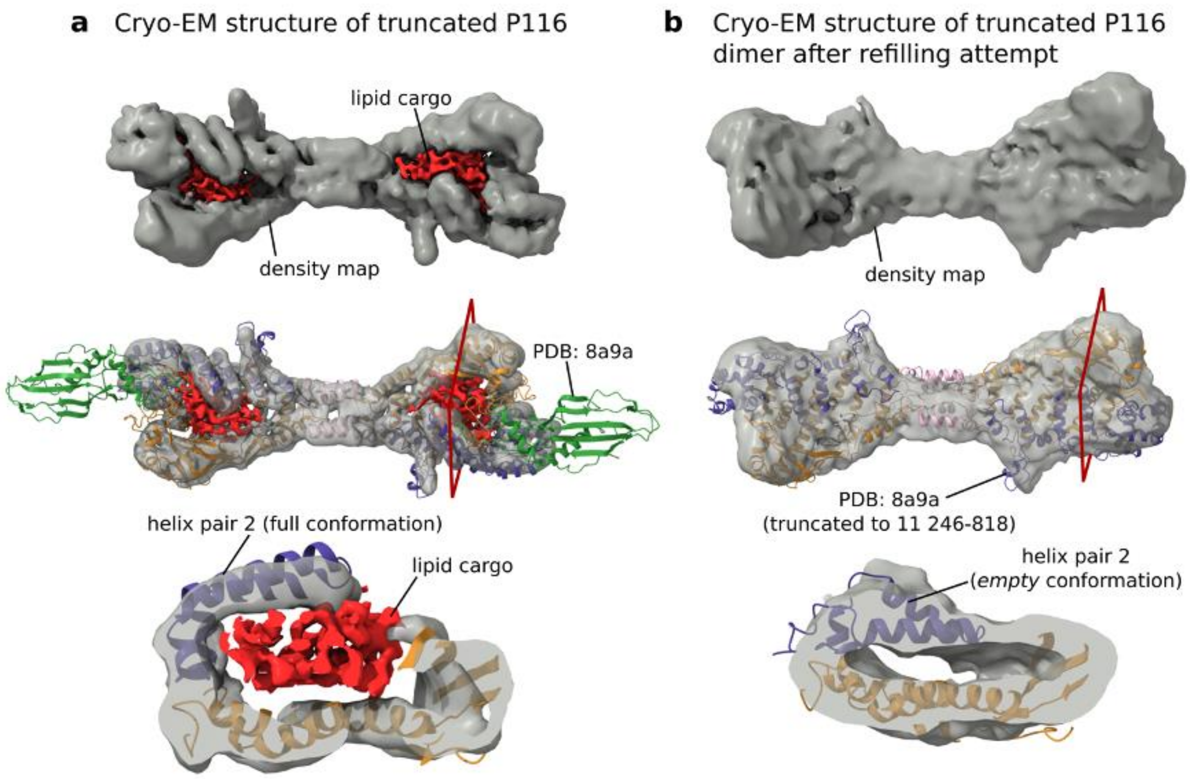

**Fig. S9. Cryo-EM structures of ΔN-P116 (246-818) before and after the lipid extraction experiment.** (**A**) **Top.** Isosurface representation of cryo-EM density map of ΔN-P116 (246-818) in dark grey at 4.3 Å resolution (EMD-50314, PDB: 9FCH) Supplemental Figure 5a). Lipid densities inside the cavity are shown in red. **Middle.** Cryo-EM density map of ΔN-P116 superimposed with a ribbon model of the *full* (PDB: 8A9A) conformation. **Bottom.** Computational slice through the cavity of P116. The position of the computational slice with regard to the density map is indicated by the red frame. ΔN-P116 is in the *full* conformation, as determined by the large volume of the cavity, the position of helix pair 2 and the presence of lipids (compare Supplemental Figure 1b). (**B**) **Top.** Isosurface representation of cryo-EM density map of the ΔN-P116 (246-818) dimer after emptying and attempted refilling in dark grey at 6.52 Å resolution (EMD-51410, Supplemental Figure 5c). **Middle.** Cryo-EM density map of ΔN-P116 after emptying and attempted refilling superimposed with ribbon model of the *empty* (PDB: 8A9B) conformation. **Bottom.** Computational slice through the cavity of P116. The red frame indicates the position of the computational slice with regard to the density map. ΔN-P116 is in the *empty* conformation, as determined by the small volume of the cavity, the position of helix pair 2and the absence of lipids (compare Supplemental Figure 1b).

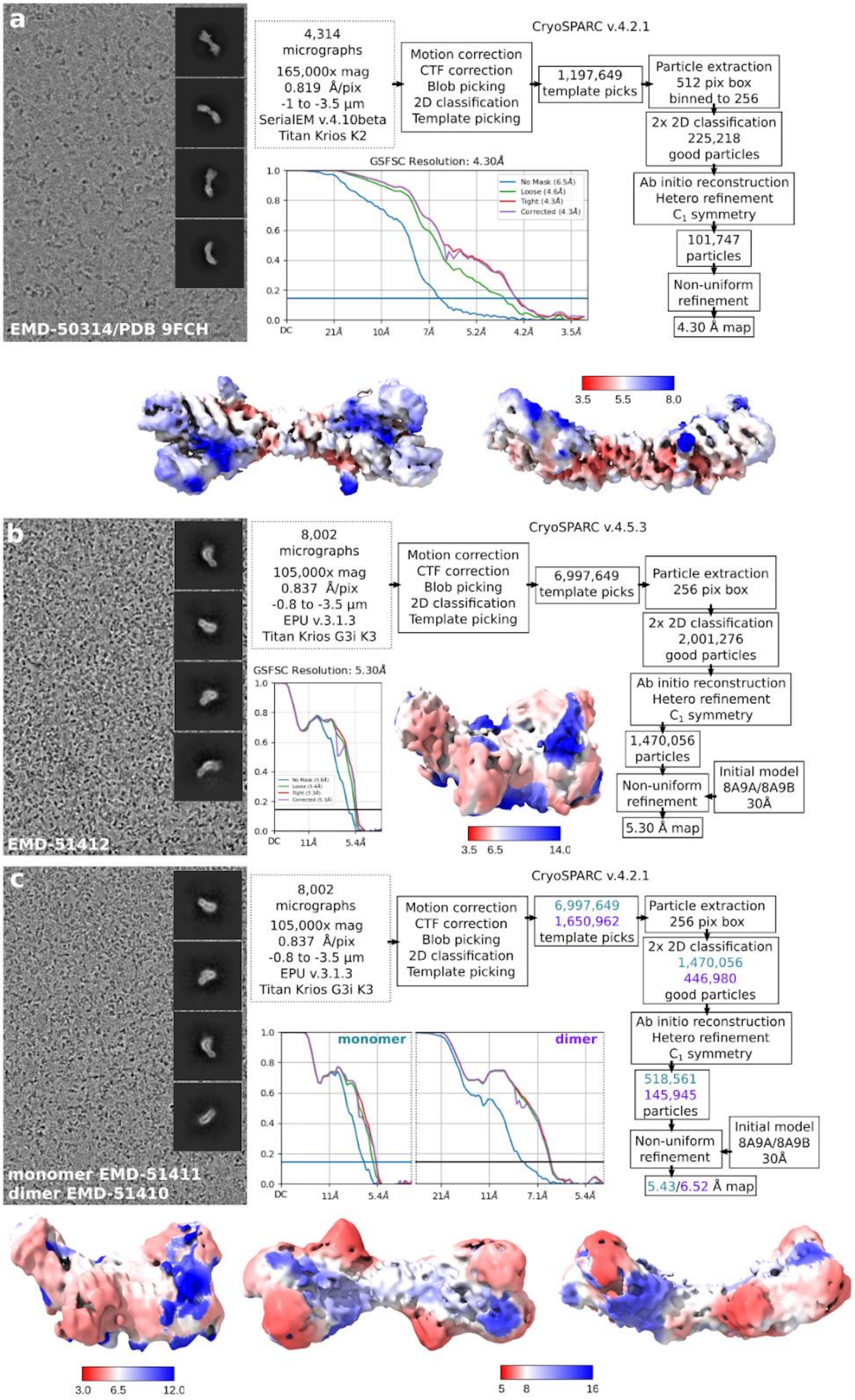
a
EMD-50314/PDB 9FCH
4,314 micrographs
165,000x mag
0.819 Å/pix
-1 to -3.5 µm
SerialEM v.4.10beta
Titan Krios K2
CryoSPARC v.4.2.1
Motion correction
CTF correction
Blob picking
2D classification
Template picking
1,197,649 template picks
Particle extraction
512 pix box
binned to 256
2x 2D classification
225,218
good particles
Ab initio reconstruction
Hetero refinement
C1 symmetry
101,747 particles
Non-uniform refinement
4.30 Å map
GSFSC Resolution: 4.30Å
3.5 5.5 8.0
b
EMD-51412
8,002 micrographs
105,000x mag
0.837 Å/pix
-0.8 to -3.5 µm
EPU v.3.1.3
Titan Krios G3i K3
CryoSPARC v.4.5.3
6,997,649 template picks
Particle extraction
256 pix box
2x 2D classification
2,001,276
good particles
1,470,056 particles
Initial model
8A9A/8A9B
30Å
5.30 Å map
GSFSC Resolution: 5.30Å
3.5 6.5 14.0
c
monomer EMD-51411
dimer EMD-51410
CryoSPARC v.4.2.1
6,997,649
1,650,962
template picks
1,470,056
446,980
good particles
518,561
145,945
particles
5.43/6.52 Å map
monomer
dimer
3.0 6.5 12.0
5 8 16

**Fig. S10. Cryo-EM data acquisition and processing.** (**A**) ΔN-P116 (residues 246–818) in the *full* conformation after purification from *E.coli*. The density map is deposited under EMD-50314/PDB: 9FCH. (**B**) Monomer of ΔN-P116 (residues 246–818) in the *empty* conformation after emptying. The density map is deposited under EMD-51412. (**C**) Monomer and dimer of ΔN-P116 (residues 246–818) in the *empty* conformation after incubation with DPPC liposomes. The density maps are deposited under EMD-51411 and EMD-51410.

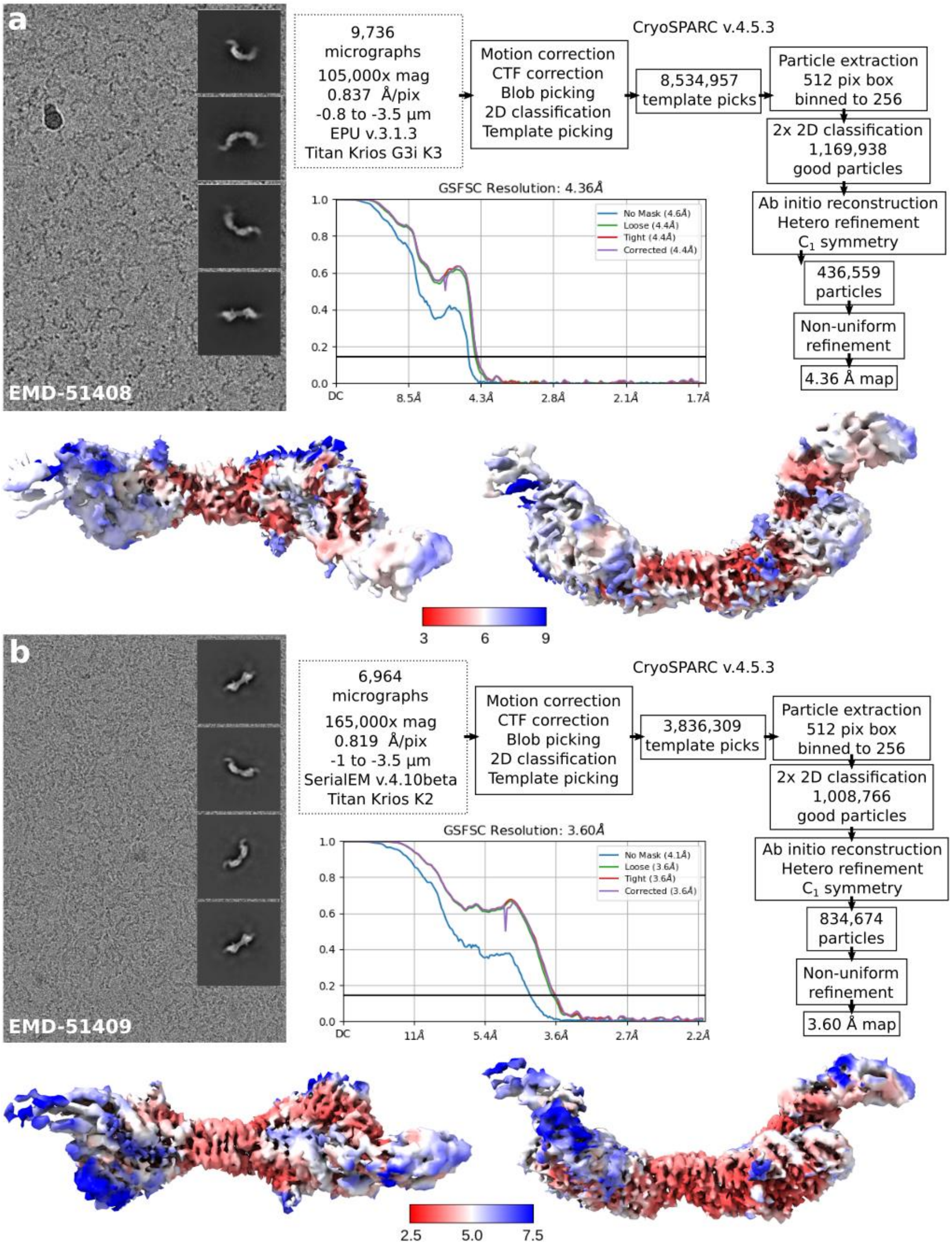

**Fig. S11. Cryo-EM data acquisition and processing.** (**A**) Full-length P116 (residues 30–957) in the *empty* conformation after emptying. The density map is deposited under EMD-51408. (**B**) Full-length P116 (residues 30–957) in the *full* conformation after incubation with DPPC liposomes. The density map is deposited under EMD-51409.

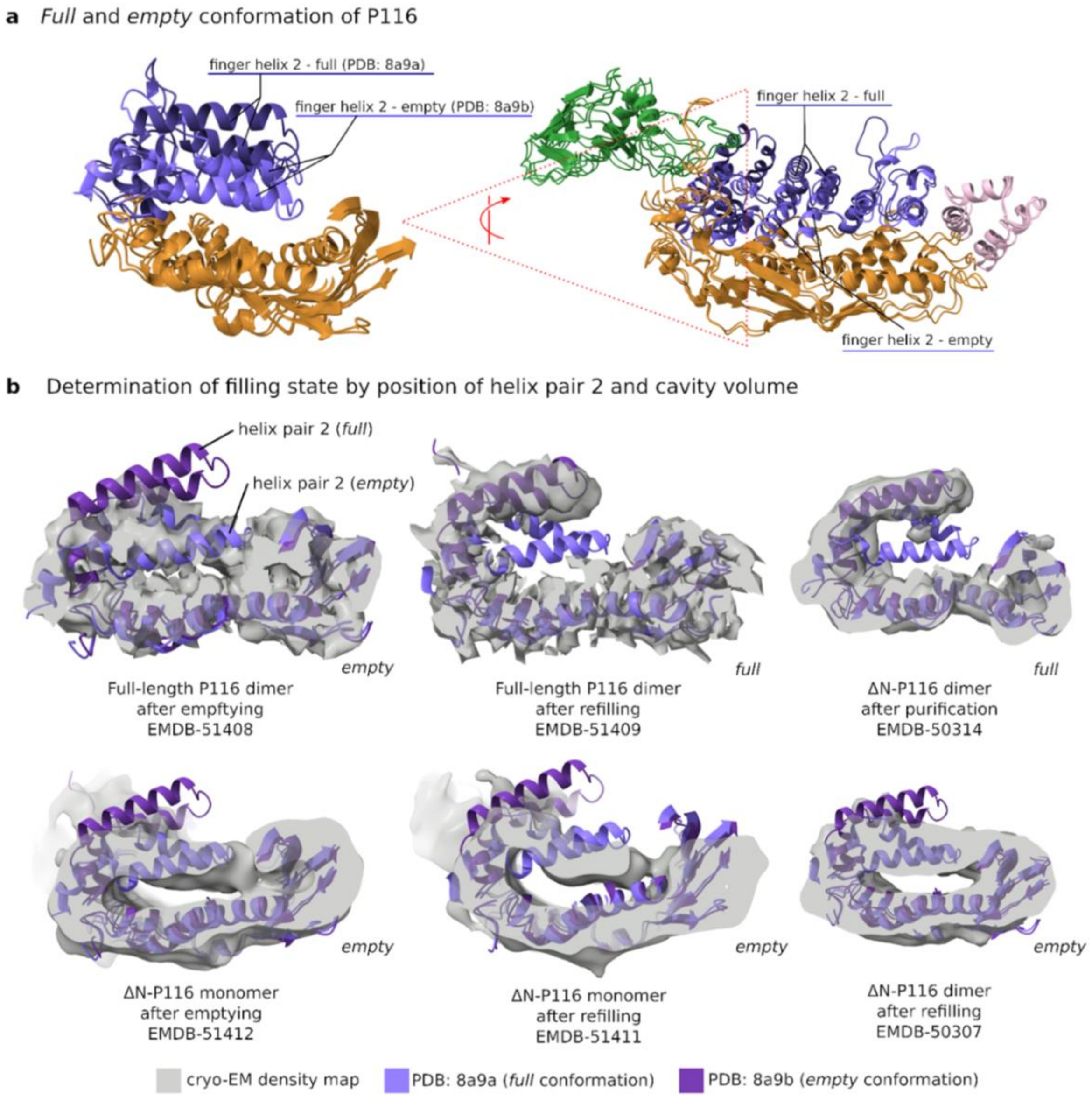

**Fig. S12. The conformation of P116 correlates with its filling state**
(**A**) Conformational comparison of *full* P116 (PDB: 8a9a) and *empty* P116 (PDB: 8a9b). *Full* and *empty* P116 are distinguished by a reduction of the hydrophobic cavity volume by 60 percent and a movement of the finger helix pairs towards the palm, most noticeably of finger helix 2 by 13 Å. The view on the right corresponds to a vertical slice through the view on the right, clipped in the position of the vertical red line. (**B**) Superposition of cryo-EM density maps of P116 from Figure 4 with ribbon models of the *full* (PDB: 8A9A, dark purple) and *empty* (PDB: 8A9B, light purple) conformation. The filling state is determined by the position of helix pair 2 and the cavity volume.

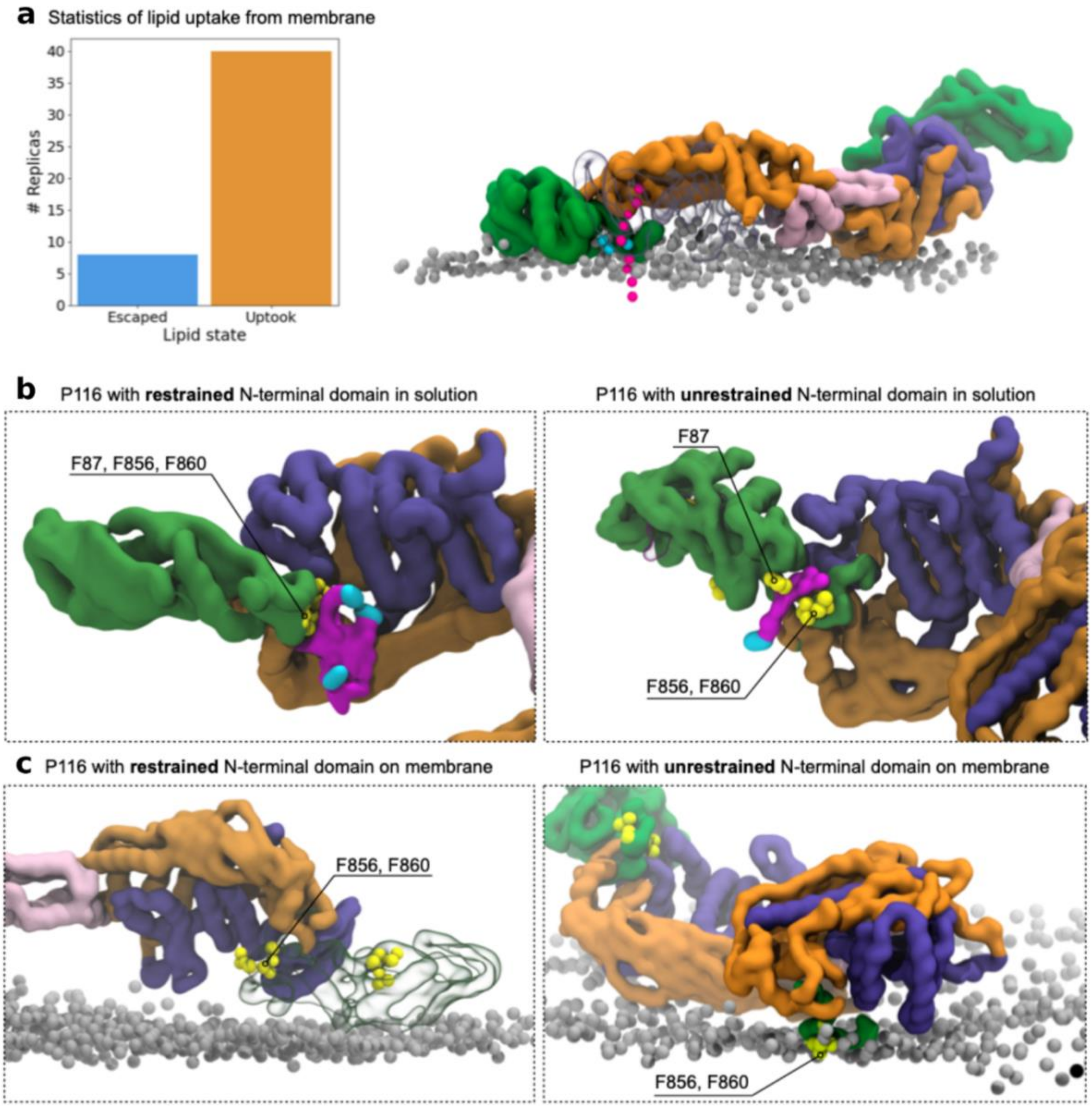

**Fig. S13. MD simulation with restrained N-terminal domain and lipid uptake statistics.** P116 (60–868) is represented by a filled volume coloured by domain. When present, the membrane is indicated by the phosphate moiety (represented by silver beads), and solvated lipids are represented by phosphate moieties (blue) and tails (purple). Water and ions are not shown for clarity. (**A**) Statistics of lipid uptake from the membrane when a lipid in the observed split position was placed in the gap between the membrane and the DCA of an N-docked P116 (as depicted in the render on the right). P116 took up the lipid in 40 of a total of 48 replicas of this system. (**B,C**) Comparison between MD simulations of P116 (60-868) with restricted versus flexible N-terminal domain. (**B**) Only the flexible N-terminal domain facilitated the caging of the lipid below the DCA and, subsequently, lipid uptake through the DCA. The constraint N-terminal domain did not allow lipid uptake. (**C**) The flexible N-terminal domain allowed the insertion of phenylalanine residues F856 and F860 into the membrane, while the constrained N-terminal domain did not show stable membrane docking.

**a** Mass spectra of peptide binding to P116

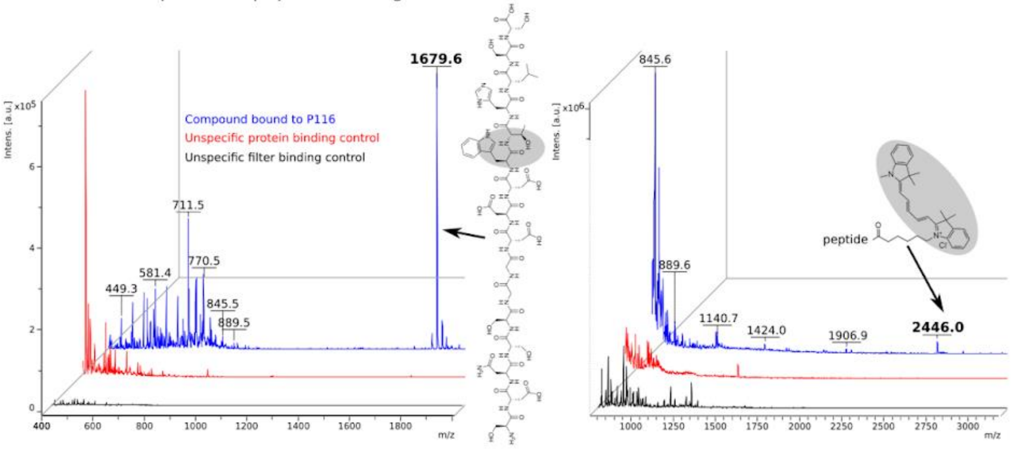

**b** Single-particle cryo-EM processing pipeline for peptide-filled P116

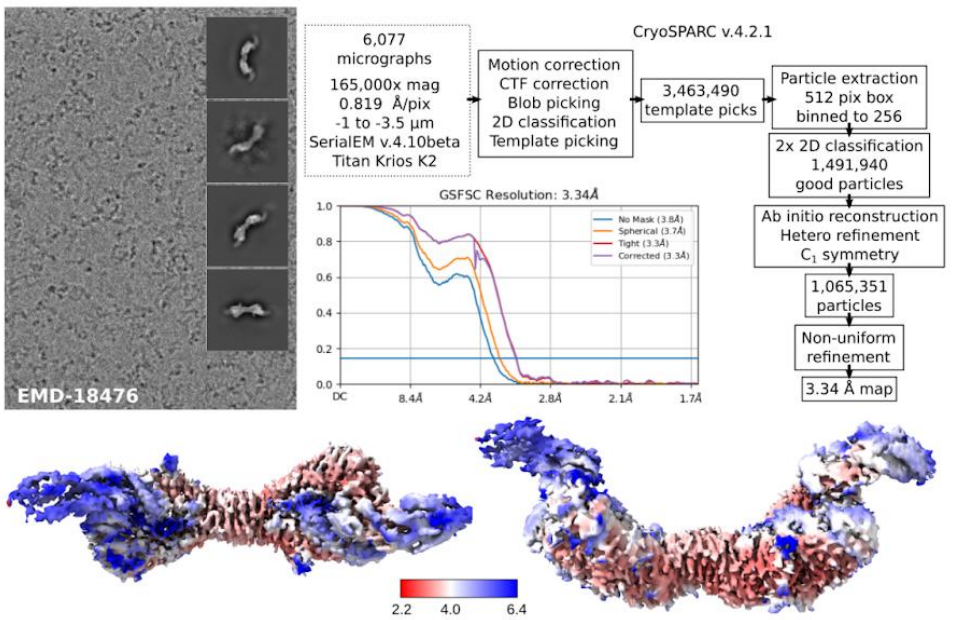

**c** Cryo-EM structure of P116 binding a peptide inside its cavity

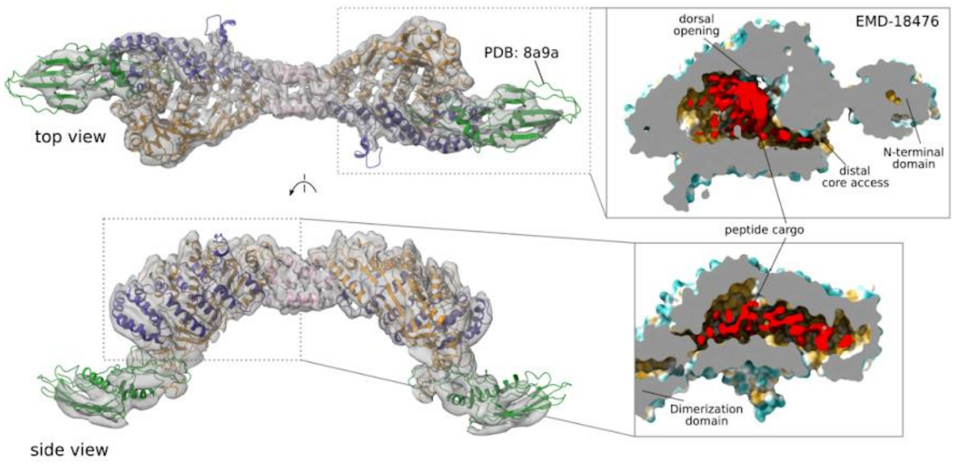

**Fig. S14. Analysis of hydrophobic peptide binding by P116.** (**A**) Mass spectra show that P116 specifically binds the plain peptide. Peptide bound to P116 (blue), to P110 or P140 (other mycoplasma membrane proteins, red) or to the filter (black). The structure of the compound is shown next to their corresponding mass peaks. The area of the compound with the largest accessible surface diameter is highlighted in grey. In the left peptide, this is the tryptophan residue (0.85 nm); in the right peptide, this is the Cy5-fluorophore (1.1 nm). (**B**) Cryo-EM processing pipeline for peptide-filled P116 (residues 30–957). The density map is deposited under EMD-18476. (**C**) Single-particle cryo-EM structure of P116 filled with the plain hydrophobic peptide. The grey surface shows the density of peptide-filled P116 at 3.3 Å. It was fitted with the structure of full P116 (colored ribbon model, PDB: 8A9A). The surface representation in the lower panels is colored by the hydrophobicity factor (yellow is hydrophobic, and blue is hydrophilic) with the sectioning surface in grey. The peptide inside the hydrophobic cavity of P116 is colored red.

**Table S1. Fragments of P116 used in this study**

| name | amino acids | missing | rationale | experiments |
|---|---|---|---|---|
| Full-length ectodomain/ full-length P116 | 30 – 975 | TMH C-terminal fragment | ectodomain lacking unstructured part of the C-terminus to facilitate expression | lipid binding and delivery assays, cryo-EM |
| ΔN-ectodomain/ ΔN-P116 | 246 – 818 | TMH, linker, N-terminal domain, C-terminal fragment | ectodomain additionally missing linker and N-terminal domain | lipid binding assays, cryo-EM |
| PDB structure of ectodomain | 60 – 868 | TMH, linker, C-terminal fragment | ectodomain missing segments that could not be resolved by cryo-EM and are thus absent in the protein PDBs | MD simulations |
| Native P116 | 1 – 1030 | - | - | Cryo-ET |

**Table S2. Statistical analysis of liposome fluorescence**

| | 1 | 2 | 3 | 4 | 5 |
|---|---|---|---|---|---|
| | Auto-fluorescence control | Unspecific transfer control with P110/140 | Lipid transfer with P116 | | |
| | | | *DPPC* | *DPPC* | DOPC |
| | | | *ultrafiltration* | *ultrafiltration + Proteinase K* | *ultrafiltration* |
| AVERAGE | 9.55 | 15.45 | 53.33 | 48.78 | 59.78 |
| STDEV | 3.34 | 5.42 | 15.17 | 16.14 | 15.29 |
| n | 21 | 21 | 44 | 44 | 44 |

| | | | |
|---|---|---|---|
| P (two tail) | 1 vs. 2 | 0.000125543 | *** |
| | 2 vs. 3 | 2.82299E-21 | *** |
| | 1 vs. 3 | 4.53119E-24 | *** |
| | 3 vs. 4 | 0.18897217 | ns |
| | 4 vs. 5 | 0,04906611 | * |

*P values originate from an unpaired T-test.*

**Table S3a. Single-particle cryo-EM data collection and processing**

| | filled full-length P116 (30–957) **EMD-51409** | empty full-length P116 (30–957) **EMD-51408** |
|---|---|---|
| **Microscope** | FEI Titan Krios | FEI Titan Krios G3i |
| **Detector** | Gatan K2 Summit | Gatan K3 Summit |
| **Acquisition Software** | SerialEM v4.1.0beta | EPU v.3.1.3 |
| **Magnification** | 165,000x | 105,000x |
| **Voltage (kV)** | 300 | 300 |
| **Electron exposure ($e^-$/Å$^2$)** | 50 | 50 |
| **Defocus range (µm)** | -0.8 to -3.5 | -0.8 to -3.5 |
| **Pixel size (Å)** | 0.819 | 0.837 |
| **Symmetry imposed** | C1 | C1 |
| **Initial particle images** | 3,836,309 | 8,534,957 |
| **Final particle images** | 834,674 | 436,559 |
| **Map Resolution (Å)** | 3.60 | 4.36 |
| **Map Resolution (Å) EMDB** | 4.07 | 4.64 |
| **FSC threshold** | 0.143 | 0.143 |
| **Map resolution range (Å)** | 2.4 - 9 | 2.8 – 9.6 |
| **Number of frames** | 50 | 50 |
| **Micrographs used** | 6,964 | 9,736 |
| **Processing software** | cryoSPARC v4.5.3 | cryoSPARC v4.5.3 |
| **Motion correction** | cryoSPARC v4.5.3 | cryoSPARC v4.5.3 |
| **CTF estimation** | cryoSPARC v4.5.3 | cryoSPARC v4.5.3 |
| **Particle images after 2D classification** | 1,008,766 | 1,169,938 |
| **Map sharpening B factor** | -124.99 | -180.25 |

**Table S3b. Single-particle cryo-EM data collection and processing**

| | filled truncated P116 (246–818)<br><br>**EMD-50314**<br>**PDB-9FCH** | empty monomer of ΔN-P116 (246–818)<br><br>**EMD-51412** | empty monomer of ΔN-P116 (246–818) after refilling attempt<br>**EMD-51411** |
|---|---|---|---|
| **Microscope** | FEI Titan Krios | FEI Titan Krios G3i | FEI Titan Krios G3i |
| **Detector** | Gatan K2 Summit | Gatan K3 Summit | Gatan K3 Summit |
| **Acquisition Software** | SerialEM 4.10beta | EPU v.3.1.3 | EPU v.3.1.3 |
| **Magnification** | 165,000x | 105,000x | 105,000x |
| **Voltage (kV)** | 300 | 300 | 300 |
| **Electron exposure ($e^-$/Å$^2$)** | 50 | 50 | 50 |
| **Defocus range (µm)** | -1 to -3.5 | -0.8 to -3.5 | -0.8 to -3.5 |
| **Pixel size (Å)** | 0.819 | 0.837 | 0.837 |
| **Symmetry imposed** | C1 | C1 | C1 |
| **Initial particle images** | 1,197,649 | 6,997,649 | 4,386,901 |
| **Final particle images** | 101,747 | 1,470,056 | 518,561 |
| **Map Resolution (Å) cryoSPARC** | 4.30 | 5.30 | 5.43 |
| **Map Resolution (Å) EMDB** | 6.5 | 5.78 | 3.62 |
| **FSC threshold** | 0.143 | 0.143 | 0.143 |
| **Map resolution range (Å)** | 3.5–9 | 4-12 | 4–13 |
| **Number of frames** | 50 | 50 | 50 |
| **Micrographs used** | 4,314 | 8,002 | 6,128 |
| **Processing software** | cryoSPARC v4.2.1 | cryoSPARC v4.5.3 | cryoSPARC v4.2.1 |
| **Motion correction** | cryoSPARC v4.2.1 | cryoSPARC v4.5.3 | cryoSPARC v4.2.1 |
| **CTF estimation** | cryoSPARC v4.2.1 | cryoSPARC v4.5.3 | cryoSPARC v4.2.1 |
| **Particle images after 2D classification** | 225,218 | 2,001,276 | 1,470,056 |
| **Map sharpening B factor** | -175 | -354.77 | 407.9 |
| **Map without sharpening** | - | - | EMD-18478 |

**Table S3c. Model building statistics of filled truncated P116 (246–818) (EMD-50314, PDB-9FCH)**

| | |
|---|---|
| **Model composition** | |
| Non-hydrogen atoms | 4,536 |
| Protein residues | 573 |
| Ligands | 0 |
| **R.m.s. of z-score** (number of standard deviations from expected value) | |
| Bond lengths | 0.27 |
| Bond angles | 0.48 |
| **Validation** | |
| Clashscore | 11 |
| Poor rotamers (%) | 1.4 |
| **Ramachandran plot** | |
| Favored (%) | 90 |
| Allowed (%) | 10 |
| Disallowed (%) | 0 |

**Table S4. Assessment of the conformation/filling state of P116.** To determine the conformation of P116, the cryo-EM density maps at a defined contour level were rigid-body fitted (chimerax *fitmap*) with both PDB: 8A9B (*empty* conformation of P116) and PDB: 8A9A (*full* conformation of P116). Subsequently, the correlation of the map-to-map fit of helix pair 2 (aa 444 – 476) at 3 Å resolution into the density map was measured (chimerax *molmap* and chimeraX *measure correlation*).

| | | | **PDB: 8A9B** | **PDB: 8A9A** |
|---|---|---|---|---|
| | | | ***empty* conformation** | ***full* conformation** |
| **density map** | **EMDB-ID** | **contour level** | **correlation** | **correlation** |
| Full-length ectodomain<br>• after emptying<br>• dimer | 51408 | 0.0746 | 0.7092 | 0.3282 |
| Full-length ectodomain<br>• after refilling<br>• dimer | 51409 | 0.18 | 0.3393 | 0.7851 |
| ΔN-P116<br>• after purification<br>• dimer | 50314 | 0.197 | 0.4899 | 0.7824 |
| ΔN-P116<br>• after emptying<br>• monomer | 51412 | 0.0392 | 0.6867 | 0.4567 |
| ΔN-P116<br>• after refilling<br>• monomer | 51411 | 0.0384 | 0.615 | 0.3342 |
| ΔN-P116<br>• after refilling<br>• dimer | 51410 | 0.112 | 0.7876 | 0.7068 |

**Table S5. Single-particle cryo-EM data collection and processing**

| | peptide filled P116 (30–957) **EMD-18476** |
|---|---|
| **Microscope** | FEI Titan Krios |
| **Detector** | Gatan K2 Summit |
| **Acquisition Software** | SerialEM 4.10beta |
| **Magnification** | 165,000x |
| **Voltage (kV)** | 300 |
| **Electron exposure ($e^{-}/Å^{2}$)** | 50 |
| **Defocus range (µm)** | -1 to -3.5 |
| **Pixel size (Å)** | 0.819 |
| **Symmetry imposed** | C1 |
| **Initial particle images** | 3,463,490 |
| **Final particle images** | 1,065,351 |
| **Map Resolution (Å)** | 3.34 |
| **FSC threshold** | 0.143 |
| **Map resolution range (Å)** | 2.5–8 |
| **Number of frames** | 22 |
| **Micrographs used** | 6,007 |
| **Processing software** | cryoSPARC v4.2.1 |
| **Motion correction** | cryoSPARC v4.2.1 |
| **CTF estimation** | cryoSPARC v4.2.1 |
| **Particle images after 2D classification** | 1,491,940 |
| **Map sharpening B factor** | -120 |

**Movie S1. MD trajectory showing P116 docking to the membrane.** P116 (60–868) is represented by a filled volume coloured by domain. the membrane is indicated by the phosphate moiety (represented by silver beads). Water and ions are not shown for clarity.

**Movie S2. MD trajectory showing the uptake of a lipid through the DCA from solvent.** P116 (60–868) is represented by a filled volume coloured by domain. The acquired lipid is represented with the phosphate moieties in cyan and the tails in pink. The other lipids are represented by silver beads. Water and ions are not shown for clarity.
